# Deciphering AMP deaminase-2 structure, activators and regulators underpinning cellular function in human fructose and nucleotide metabolism

**DOI:** 10.64898/2026.06.10.731346

**Authors:** Ashley M. Rebelo, Nemanja Vuksanovic, Lanlan Han, Dean R. Tolan, Karen N. Allen

## Abstract

AMP deaminase (AMPD) plays an integral role in fructose metabolism *via* its regulation by ATP, GTP and phosphate (Pi). The fructose catabolic pathway consumes ATP, producing ADP, which is further metabolized to AMP, triggering a cascade of reactions initiated by AMPD. This degradative pathway results in the final product uric acid, which is associated with metabolic acidosis, mitochondrial dysfunction, and gout. Understanding the regulation of the human liver AMPD isozyme (hAMPD2-2) under physiological conditions and under fructose consumption conditions will enable the design of targeted therapeutics to block the accumulation of uric acid. We report the first successful expression and purification from *Escherichia coli* of both the full-length and catalytic domains of hAMPD2-2. Steady-state kinetics confirmed allosteric activation by ATP of both the full-length and catalytic domains of hAMPD2-2 at physiological ATP concentrations (2-5 mM), suggesting that the allosteric ATP-binding site is located in the catalytic domain. Competitive inhibition by GTP of the ATP-activated enzyme, with K_i_ values of 74 and 101 μM for the full-length and catalytic domains, respectively, was also consistent with this regulatory model. P_i_, previously described in yeast AMPD as a competitive inhibitor, was shown to play a more nuanced role, that of enhancing inhibition of hAMPD2-2 when the enzyme is complexed to GTP, *via* competition at the ATP allosteric site. P_i_ binding thus further inhibits the pathway under normal physiological conditions, limiting production of cellular uric acid unless and until P_i_ and GTP levels are low.

## INTRODUCTION

The United States is currently the largest sugar consumer in the world, with an average of 126.4 grams of sugar consumed per capita daily.^1^ This represents 300% of the daily recommended amount of added sugar for individuals according to FDA guidelines,^2, 3^ a result of easy access and low-cost production of this food additive. Sugar is ubiquitous in foods- in fruits, as a sweetener and in many processed foodstuffs. This universal additive is especially problematic for individuals with hereditary fructose intolerance (HFI), an autosomal recessive disease that affects 1 in 20,000 people worldwide.^4^ These patients are unable to metabolize fructose due to mutations in the ALDOB gene.^5, 6^ Typically, once fructose enters the liver (**Figure 1**), it is phosphorylated to fructose 1-phosphate (F1P) and cleaved by aldolase B. In HFI, mutations in the ALDOB gene diminish or obviate the cleavage of F1P, which accumulates, leading to liver and kidney toxicity, and possibly resulting in death.^7^ Excessive fructose ingestion also has a detrimental effect in healthy individuals.^8^ Although F1P accumulates only transiently,^9^ the rapid ATP depletion and sequestration of P_i_ in F1P activate liver AMPD, resulting in a transient acute uric acidemia.^7^ AMP is provided by converting the ADP from fructose phosphorylation to AMP by the action of adenylate kinase, which is irreversibly deaminated by AMP deaminase (AMPD2) into IMP and ammonia (**Figure 1**), leading to the final product, uric acid. Repeated boluses of accumulated uric acid are thought to be a contributor to many chronic diseases, such as gout and metabolic syndrome.^7, 10^

**Figure 1.**
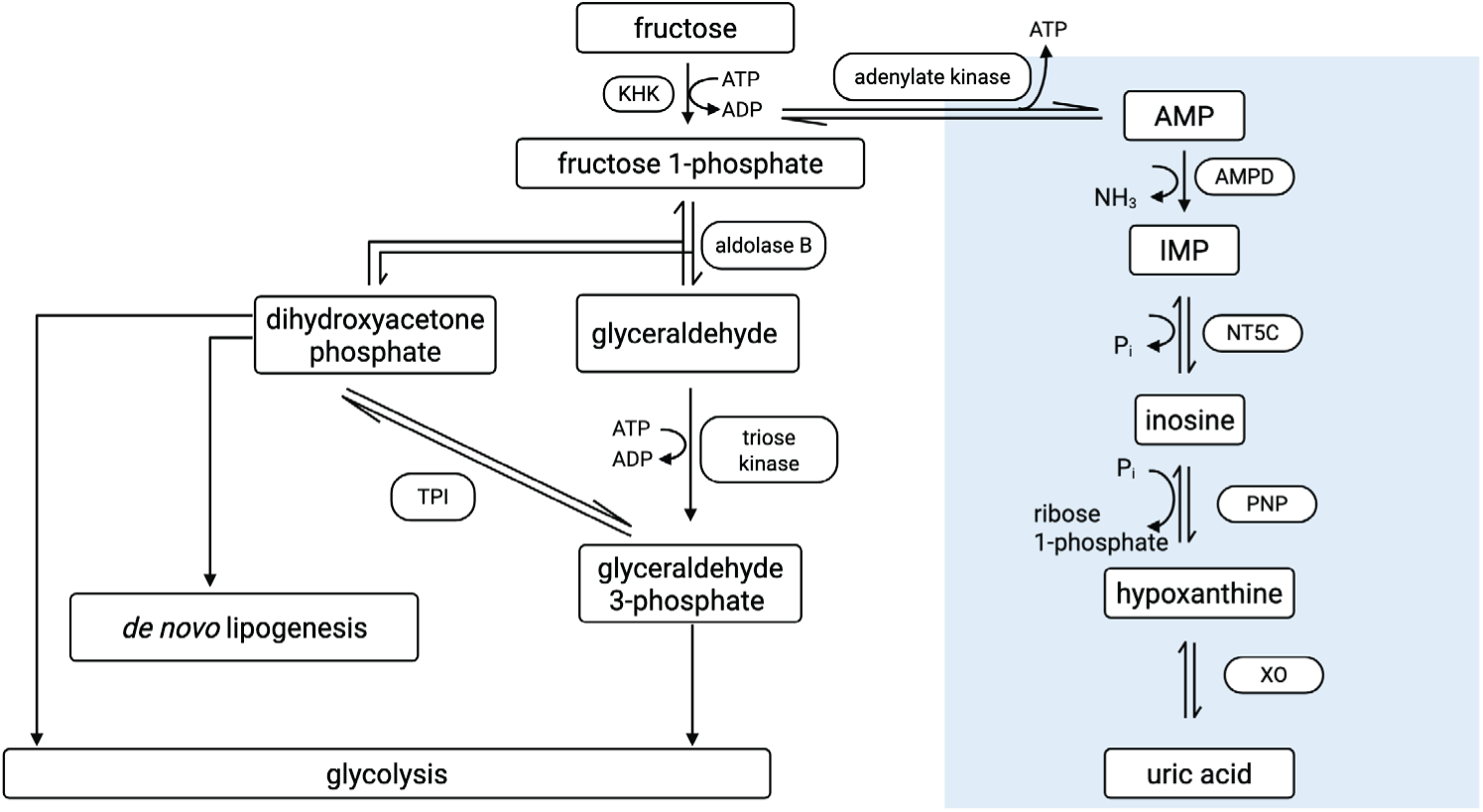
Liver fructose metabolism. n HFI, fructose 1-phosphate and uric acid concentrations increase upon fructose consumption, leading to liver and kidney failure (TPI, triose phosphatase isomerase; IMP, inosine 5’-monophosphate; P_i_, inorganic phosphate; NT5C, cytosolic 5’-nucleosidase; PNP, purine nucleoside/inosine phosphorylase; XO, xanthine oxidase). The blue square denotes the AMP catabolic pathway. Figure generated in BioRender.

AMPD plays a part in both the purine and fructose metabolic pathways. There are three isoforms of human AMPD, based on primary location,^11^ hAMPD1 (muscle and brain), hAMPD2 (liver) and hAMPD3 (erythrocytes), with hAMPD1 and hAMPD3-1 having 55.3% and 51.3% sequence identity compared to hAMPD2-2, respectively. AMPD is comprised of two domains, a short N-terminal domain (∼ 86-128 residues), previously described as regulatory,^11, 12^ and a large C-terminal catalytic domain. The C-terminal domain includes highly conserved catalytic residues, whereas the isoforms differ greatly in the sequence identity of their N-terminal domain (< 26% sequence identity between N-terminal domains, see **Figure S1A**).^12, 13^ Moreover, there are three isoforms of hAMPD2 comprised of the catalytic domain (**Figure S1B**) and several different alternatively spliced sequences encoding the N-terminal domain: hAMPDA2-1B/2 (2-2, Uniprot Q01433-2), hAMPDA2-1A/2 (2-4, Uniprot Q01433-4), hAMPDA2-1B/3 (2-5, Uniprot Q01433-5).^14^ The hAMPD2-1B/2, also known as hAMPD2-2, has the longest N-terminal domain (128 amino-acid residues). The N-terminal domain has been hypothesized to regulate enzyme activity because only this segment differs among isoforms.^12, 14^ and comparative studies of full-length and proteolyzed yeast AMPD suggest a 100-fold increase in enzyme turnover rate upon proteolysis of the N-terminal domain.^15^ As hAMPD2-2 catalyzes the first committed step leading to uric acid production (**Figure 1**), its regulation is essential in keeping ATP levels high and intracellular uric acid levels low under normal physiological conditions. Molecular-level knowledge of liver hAMPD regulation is critical to understanding the interplay between fructose metabolism and the action of this enzyme in both diseased and healthy individuals.

There are two regulators with opposing effects on AMPD2: ATP, an activator, and GTP and P_i_, both inhibitors.^15, 16^ Previous work showing ATP activation and GTP and P_i_ inhibition of yeast AMPD was, for the most part, conducted on the catalytic domain due to the instability of the full-length enzyme.^15, 17^ In addition, studies on the three isoforms of hAMPD suggested a regulatory role of the small N-terminal domain in ATP activation,12,15 pointing to a potential ATP-binding site in this domain. Herein, we present steady-state kinetic analysis of the liver hAMPD2-2 isozyme (the longest isoform) to determine the effect of ATP, GTP, and P_i_ regulation on the full-length enzyme and the C-terminal catalytic domain. Our findings show that the presence of the N-terminal domain does not influence the effect of these molecules on catalysis, therefore localizing the allosteric site(s) to the C-terminal catalytic domain. Deciphering the competitive versus non-competitive nature of the effects of these metabolites sheds light on the regulation of hAMPD2-2 under normal physiological conditions, and on how the three regulators interact with enzyme function.

## MATERIALS AND METHODS

### His_6_-tagged hAMPD2-2 expression

His_6_-tagged hAMPD2-2 gene (Uniprot ID Q01433-2) in pET28a vector (a gift from Joshua D. Rabinowitz, Princeton University) was transformed into *E. coli* BL21(DE3) cells (New England Biolabs), and colonies were selected on LB-agar plates containing 50 μg/mL kanamycin. A 250 mL overnight culture in Miller’s Luria broth with kanamycin was seeded with a single colony and grown overnight at 37 °C. Large scale (500 mL) cultures of autoinduction media (6 g tryptone, 12 g yeast extract, 25 mM Na_2_HPO_4_, 25 mM KH_2_PO_4_, 50 mM NH_4_Cl, 5 mM Na_2_SO_4_, 20 mM MgSO_4_, 0.5% glycerol, 0.2% *α*-D-lactose, 0.05% glucose, 100 μL of trace metals (Teknova)) containing kanamycin (0.15 mg/mL) were inoculated using a 1:50 dilution of the starter culture and grown at 37 °C to OD_600_ = 0.7. The temperature was lowered to 16 °C and the culture grown 14-16 hours. Cells were harvested at 5,383 x g for 20 minutes at 4 °C.

### His_6_-SUMO-hAMPD2-2 cloning and expression

DNA encoding hAMPD2-2 from pET28a described above was ligated into a pTB146-SUMO vector from the pET28a-His_6_-hAMPD2-2 plasmid using SalI and BamHI restriction enzymes. pTB146 His_6_-SUMO-hAMPD2-2 was transformed into BL21 (DE3) competent cells with pTf16 chaperone plasmid (Takara Bio), and colonies were selected on LB-agar plates containing 100 μg/mL ampicillin and 20 μg/mL chloramphenicol. A 250 mL overnight culture in Miller’s Luria broth containing 100 μg/mL ampicillin and 20 μg/mL chloramphenicol was seeded from a single colony and grown at 37 °C overnight. For large-scale production, 1 L cultures containing ampicillin, chloramphenicol and L-arabinose (0.5 mg/mL) were inoculated using a 1:50 dilution and grown at 37 °C until OD_600_ = 0.5-0.6, followed by a two-hour incubation at 30 °C before induction with 0.4 mM isopropyl IPTG and 14-16 hours growth at 16 °C. Cells were harvested at 5,383 x g for 20 minutes.

### Construction and expression of His_6_-SUMO-hAMPD2-2^D128^

Removal of the first 128 residues of the hAMPD2-2 sequence from the pET28a vector encoding His_6_-tagged hAMPD2-2 was performed *via* PCR amplification using Taq polymerase (Invitrogen), forward primer (Eurofins) containing a NdeI restriction site (5’-GACT**CATATG**CGGGAGCGTGATGTGCTG-3’) and reverse primer (Eurofins) with a HindIII restriction site (5’- CCGC**AAGCTT**GTCGACTTATTG-3’). The amplified and purified PCR product and pET28a-hAMPD2-2 were digested in NEB buffer 2™ using NdeI and HindIII restriction enzymes (New England Biolabs) and ligated using T4 ligase buffer and T4 ligase enzyme (New England Biolabs) at 16 °C overnight. Ligated pET28a-hAMPD2-2^Δ128^ was transformed into NEB5α competent cells (New England Biolabs), extracted and the sequence confirmed *via* Sanger sequencing (Genewiz). A SalI restriction site (5’-GACT**GTCGA**CGGGAGCGTGATGTGCTG-3’) was added to the sequence before the start of the catalytic domain sequence, and a BamHI restriction site (5’-GACT**GGATCC**TTATTGAGGCCCTGGGCTC-3’) was added to the 3’-end of the hAMPD2-2 sequence after the stop codon for ligation into a SalI/BamHI digested pTB146 SUMO vector. The resulting pTB146 including the hAMPD2-2^Δ128^ sequence was transformed into competent DH5*α E. coli* cells (New England Biolabs) for DNA extraction, and successful ligation confirmed by Sanger sequencing (Genewiz).

pTB146 His_6_-SUMO-hAMPD2-2^Δ128^ was transformed into BL21(DE3) competent *E. coli* cells (C2527H, New England Biolabs) together with the pG-Tf2 chaperone plasmid (Takara Bio) and colonies were selected on Miller’s Luria broth agar plates including 100 μg/mL ampicillin and 20 μg/mL chloramphenicol. A 250 mL overnight culture in Miller’s Luria broth containing ampicillin and chloramphenicol was seeded by a single colony and grown overnight at 37 °C. Large-scale 1 L cultures containing 100 μg/mL ampicillin, 20 μg/mL chloramphenicol and tetracyclin (5 ng/mL) were inoculated using a 1:50 dilution and grown at 37 °C until OD_600_ reached 0.5-0.6, followed by a 2-hour incubation at 30 °C before induction with 0.4 mM IPTG and 14-16 hours growth at 16 °C. Cells were harvested using a Beckman Avanti J-E centrifuge, JLA-9.1 rotor at 5,383 x g for 20 minutes.

### His_6_-tagged hAMPD2-2 purification

All steps were performed at 4 °C. Four 1-L pellets were combined, resuspended with lysis buffer (25 mM HEPPS pH 8, 0.5 M KCl, 50 μM zinc acetate, 10 mM imidazole, 6 mg/100 mL resuspension of DNase I (GoldBio), EDTA-free protease inhibitor (Thermofisher), 0.1% beta-mercaptoethanol (BME), 1 mM ATP, 0.25 M TMAO, 0.25 M sarcosine), and microfluidized at 18,000 psi. Lysed cells were centrifuged in a Beckman Coulter Optima L-100K ultracentrifuge at 106,255xg for 35 minutes at 4 °C. Ammonium sulfate was added to the soluble fraction to reach 40% saturation, the solution was homogenized and centrifuged at 10,000 x g, 4 °C for 10 minutes. The precipitant was resuspended in 25 mM HEPPS pH 8, 0.5 M KCl, 10 mM imidazole, 10% glycerol and 50 μM zinc acetate, and dialyzed at 4 °C for 2 hours against the same buffer to remove ammonium sulfate. The protein solution was purified on an AKTA Pure using a HisTrap pre-packed Ni affinity column with a loading buffer comprised of 25 mM HEPPS pH 8, 0.5 M KCl, 50 μM zinc acetate, 10 mM imidazole. After sample application, the column was washed with 20 CV of loading buffer. The protein was eluted from the column using elution buffer, comprised of 25 mM HEPPS pH 8, 0.5 M KCl, 50 μM zinc acetate, 250 mM imidazole by a three-step gradient: 5 CV of 0.2 x elution buffer, 5 CV of 0.3x elution buffer and 7 CV of 1x elution buffer. The protein was concentrated from the 1x elution buffer, and buffer exchanged into 25 mM HEPPS pH 8, 0.5 M KCl, 10% glycerol, 50 μM zinc acetate using an Amicon 30 kDa concentrator at 4000 x g, 4 °C. Protein concentration was determined *via* Bradford protein assay (Thermofisher) and purity was determined *via* SDS-PAGE. Samples were flash frozen and stored at -80 °C for long-term storage.

### Purification of SUMO-tagged hAMPD2-2 and hAMPD2-2^Δ^_128_

The purification of SUMO-tagged proteins was performed similarly to the full-length His-tagged enzyme purification for the steps of cell lysis and ammonium sulfate precipitation. All steps were performed at 4 °C. After dialysis to remove the excess ammonium sulfate, the protein solution was purified on an AKTA Pure using a pre-packed HisTrap affinity nickel column (Cytiva). A buffer comprised of 25 mM HEPPS pH 8, 0.5 M KCl, 50 μM zinc acetate and 25 mM imidazole was used to load the protein. The protein was eluted from the column by a four-step gradient: 5 CV of 10% elution buffer (25 mM HEPPS pH 8, 0.5 M KCl, 50 μM zinc acetate, 250 mM imidazole), 5 CV of 25% elution buffer, 6 CV of 50% elution buffer and 7 CV of 100% elution buffer. Protein was concentrated using a 30 kDa Amicon concentrator in a Beckman Avanti JS5.3 rotor at 4,000 x g at 4 °C. The purified sample was buffer-exchanged in the same buffer system as the full-length construct and characterized by Bradford protein assay (Thermofisher) and SDS-PAGE.

### Steady-state kinetic analysis of hAMPD2-2

Steady-state kinetic assays were performed using 5 nM purified His_6_-SUMO-hAMPD2-2^Δ128^ or 50 nM purified His_6_-hAMPD2-2 using the following cocktail: 50 mM TEA-HCl pH 7.4, 100 mM KCl, 4 mM MgCl_2_, 5 mM NAD^+^, 1 mM DTT, 0.05% BSA, 0.1 U/mL of cpIMPDH (1 U = 1 μmol of substrate per minute. The coupling enzyme cpIMPDH (donated by Lizbeth Hedstrom, Brandeis University) was expressed and purified by a minor modification of the published method^18^ (see SI). Assays varying AMP substrate were performed using two-fold serial dilutions from 5 mM to 39 μM. ATP activation of hAMPD2-2 assays were done by the addition of 0 – 5 mM ATP in the final assay. GTP inhibition of hAMPD2-2 assays were performed by addition of 0 – 1 mM GTP in the final assay. Assays to test P_i_ inhibition of hAMPD2-2 were done by addition of 0 – 50 mM potassium phosphate in the final assay. K_i_ values were determined using the competitive inhibition Equation (1) from GraphPad Prism, where K_Mapp_ is the apparent K_M_ of hAMPD2-2^Δ128^ with the tested compound, [I] is the compound concentration and K_i_ is the inhibition constant:

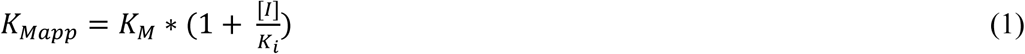

Assays varying ATP were performed at saturating substrate concentrations using two-fold serial dilutions of ATP from 5 mM to 39 μM, with 1 mM AMP in the cocktail solution. Assays varying potassium phosphate used a two-fold serial dilution from 5 mM to 39 μM, with 1 mM AMP; to this same assay, either 2 mM ATP alone or 2 mM ATP and 1 mM GTP were added to test the effect of these ligands in the presence of P_i_.

The A_340_ was read in 10-second intervals for 5 minutes using a SpectraMax M5® plate reader (Molecular Devices). The ΔA_340_ was calculated for each 10-second interval and averaged over the linear region of the raw data (constant rate), as determined by data reduction analysis (R2 > 0.99) in the SoftMax Pro software of the SpectraMax M5® plate reader (Molecular Devices). Outlier replicates were determined using Grubb’s test in GraphPad Prism and samples with a Z score > 3 or < -3 were removed from analysis. Activity was fit to the Michaelis-Menten equation using GraphPad Prism. To determine the inhibition mode, steady-state kinetics data were fit to a global mixed-model inhibition regression curve to determine α using Equation 2:

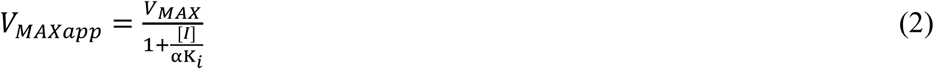

### hAMPD2 steady-state kinetics varying enzyme activator and inhibitor

Enzyme activity using saturating amounts of AMP (1 mM) with increasing amounts of ATP and varying amounts of GTP (with and without P_i_) was calculated by normalizing activity values to the activity at the highest ATP concentration (5 mM) from Equation (2). This data was fit to a four-parameter binding curve (log [agonist] vs. response in GraphPad Prism) using % activity vs log [ATP] using Equation 3:

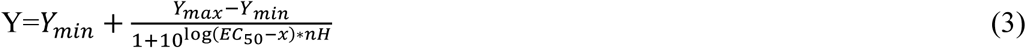

x is ATP concentration, Y is average enzyme % activity, nH is a constant describing the steepness of the curve and Y_max_ and Y_min_ are plateaus at the highest % activity and lowest % activity, respectively. EC_50_ value is the value at 50% enzyme activity at a fixed concentration of ATP.

### Assays for effect of inhibitor and activator in the same system

The hAMPD2-2^Δ128^ catalytic activity was determined using saturating substrate (1 mM AMP) with 2 mM ATP, varying P_i_ concentration with and without GTP. The fraction of enzyme inhibition (*i*) was determined using Equation 4:^19^

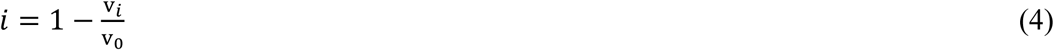

where v_i_ is the rate of the enzyme in the presence of P_i_, v_0_ is the rate of the enzyme in the absence of P_i_ (A_340_/min). Equation 5 was used to determine K_i_:

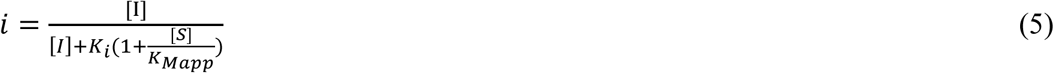

Where [I] is the concentration of P_i_, K_i_ is the inhibition constant of P_i_, [S] is the concentration of AMP, K_Mapp_ is the Michaelis-Menten constant (with and without GTP), and *i* is the fraction inhibition of enzyme.

In the absence of GTP, the rate of hAMPD2-2^Δ128^ with saturating AMP (1 mM) and varying ATP concentrations at fixed P_i_ concentrations was determined and fit to an inhibition model (Equation 6) in which inhibitor competes with the activator binding site:^19^

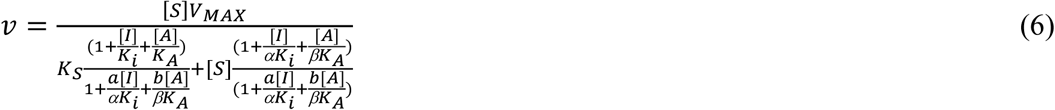

Where v is the enzyme rate, [S] is AMP concentration, V_MAX_ is the maximum enzyme rate, K_S_ is the substrate binding constant, [I] is P_i_ concentration, K_i_ is the inhibition constant of P_i_, [A] is ATP concentration, K_A_ is the binding constant of the activator, and a, b, α and β are constants.

### Global Fitting of Absorbance Traces

Absorbance traces for catalytic-domain hAMPD2-2^Δ128^ were analyzed in KinTek Explorer (v11.1.1, KinTek Corporation) using the same datasets employed for the steady-state analysis and fit using the observable A_340_ = scale·P + background, where P is the simulated product-associated species. The scale term was fixed at 0.00622 as a Beer’s law scaling factor that incorporates the NADH extinction coefficient at 340 nm and the assay path length. Experiment-specific background terms were fit as needed. The reduced kinetic model included ATP-dependent activation, AMP turnover through the ATP-activated branch, competitive inhibition by GTP, and P_i_-dependent inhibition. Fits were refined iteratively across the AMP, ATP, GTP, and P_i_ datasets, with selected association rate constants fixed to reduce overparameterization. The full reaction scheme, parameter set, and plots of the KinTek fits overlaid with the experimental data are provided in the Supporting Information.

### Nano-DSF assays

Nano-DSF assays were performed with 3.8 μM and 4.4 μM of each SUMO construct (full length and catalytic domain, respectively), diluted in 25 mM HEPPS pH 8, 0.5 M KCl, 10% glycerol, and 50 mM zinc acetate. Ligands were added at a final concentration of 2 mM. Protein was incubated with each ligand for 20 min on ice, except for the substrate AMP, which was added immediately before measurements began. Each sample was run in triplicate, and measurements were made at 330 nm and 350 nm at a ramp rate of 1 °C/min from 20 to 95 °C at 100% intensity using quartz capillaries in a Prometheus NanoDSF (NanoTemper). Melting temperatures (T_m_) were determined and averaged by location of the inflection point using the first derivative of the fluorescence at 350 nm and 330 nm.

## RESULTS

ATP is a known activator of AMPD2, but, as no structure is yet available with ATP present, the activator binding site is unknown. To better understand the regulation of AMPD2 activity and pave the way for future structural work to enable therapeutic intervention, we sought to localize the allosteric site to either the catalytic domain or the N-terminal domain, which has previously been hypothesized to confer regulation. To approach this problem, the effects of substrate, product, and activator binding on kinetics were assessed and compared between the catalytic domain alone and the full-length hAMPD2-2 isoform, which includes the N-terminal regulatory domain.

### ATP and GTP stabilize hAMPD2-2, ATP activates hAMPD2-2

The effect of ATP, GTP and P_i_ binding on hAMPD2-2 thermal stability was determined by measuring the T_m_ by nanoDSF, and comparing it to that of unliganded hAMPD2-2, and liganded to AMP substrate and IMP product. The T_m_ of unliganded full-length hAMPD2-2 is 57.4 ± 0.5 °C, and for the catalytic domain is 56.1 ± 0.4 °C (**Figure 2**). The T_m_ for both constructs increases when bound to AMP substrate (58.5 ± 0.1 °C and 60.8 ± 0.1 °C, respectively) and to a greater extent when bound to ATP (60.6 ± 0.1 °C and 62.7 ± 0.2 °C, respectively). Significant stabilization is not observed in the presence of the product IMP (58.7 ± 0.1 °C and 56.1 ± 0.2 °C, respectively) or for the catalytic domain in the presence of P_i_ (58 ± 1 °C, data not shown). The increase in T_m_ of the catalytic domain in the presence of ATP is consistent with a model in which the allosteric site is located in this domain, and not the N-terminal domain, previously proposed to play a role in regulation by ATP.^11, 12^ Addition of both ATP and GTP in a ratio of 1:1 at 2 mM each to hAMPD2-2 results in an increase in T_m_ of the unliganded enzyme by 6 and 8 °C for the full-length and catalytic domain, respectively (**Figure 2**), similar to that seen for either ligand alone. All thermal unfolding experiments on full-length and catalytic domain samples yielded a single peak. Both constructs have a SUMO tag, which did not result in a separate unfolding event during thermal unfolding (data not shown). The thermal stabilization data indicate that the ligands are not binding to the N-terminal domain of hAMPD2-2.

**Figure 2.**
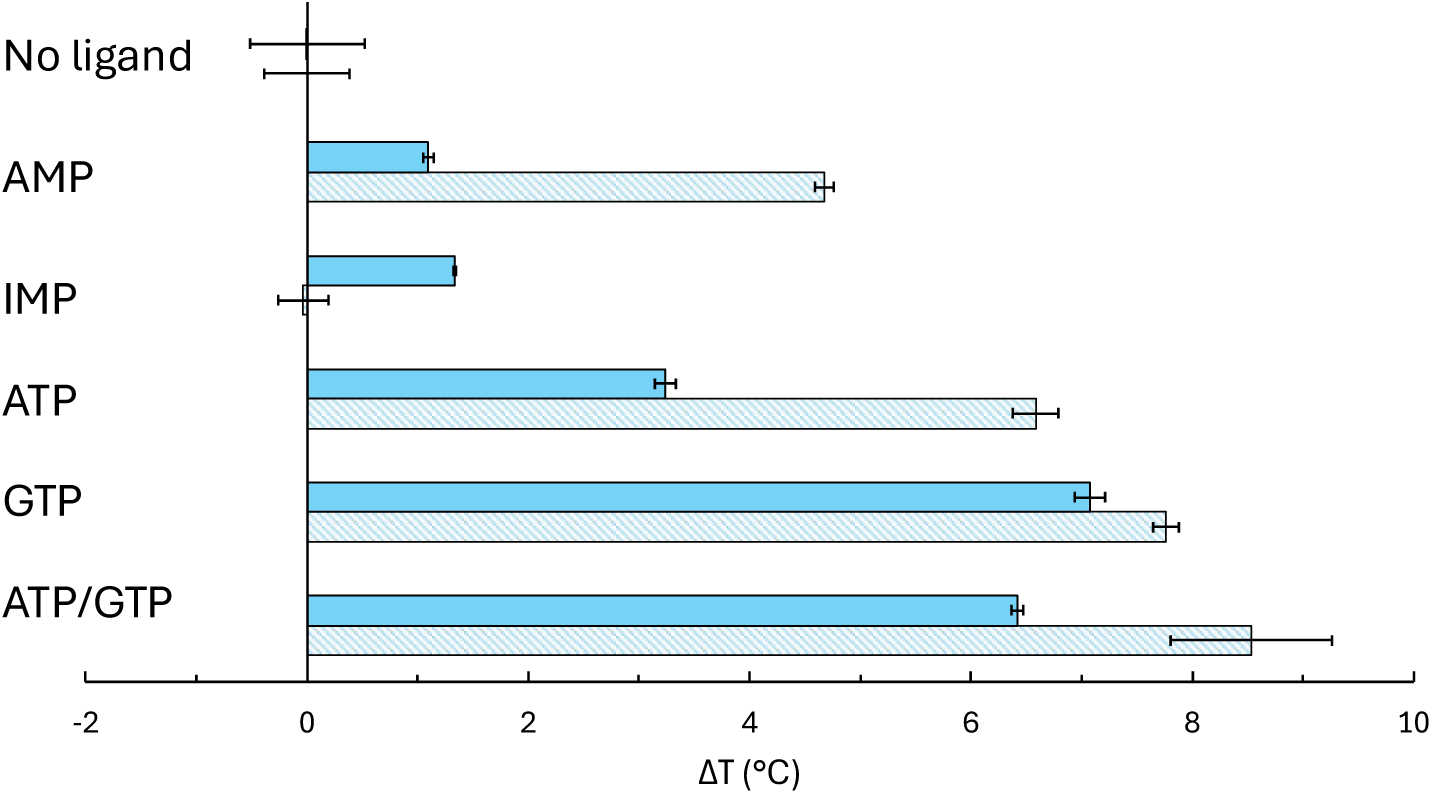
Melting temperature (T_m_) of full-length and catalytic domain of hAMPD2-2 with substrate (AMP), product (IMP), and regulators (ATP, GTP). Change in T_m_ of full-length construct (filled blue bars) and catalytic domain (striped, blue bars) in the absence and presence of 2 mM ligand. A 1:1 ratio of ATP:GTP was used at 2 mM each.

As ATP confers higher thermal stability, it was essential to determine its effect on hAMPD2-2 catalysis, with and without the N-terminal domain. Steady-state kinetics constants were determined using the Michaelis-Menten model at varying ATP concentrations. The catalytic activity of hAMPD2-2 is nearly absent without ATP, such that we could not determine steady-state kinetic constants (**Table 1**, **Figure S2A**). The enzyme is activated at ATP concentrations as low as 0.5 mM and is fully activated at 2 mM ATP, with K_M_^AMP^ = 350 ± 20 μM and k_cat_ = 13 ± 0,1 s^-1^. Similarly, for the catalytic domain of hAMPD2-2, there is limited enzyme activity in the absence of ATP, and the addition of 2 mM ATP yields full activation with a K_M_^AMP^ of = 460 ± 20 μM and k_cat_ of = 170 ± 1 s^-1^ (**Table 1**, **Figure S2B**). This data, together with that from the effect of ligands on T_m,_ supports the presence of an allosteric binding site for ATP located in the catalytic domain of hAMPD2-2. Notably, the effect of ATP is in K_m,_ whereas k_cat_ is relatively constant. At all ATP concentrations, k_cat_ is approximately ten-fold higher for the catalytic domain compared to the full-length enzyme. This was not due to unfolded protein in the full-length sample, as analyzed by DLS (data not shown).

**Table 1.** Steady-state kinetic constants at varying ATP concentrations. Values are means ± standard deviation from three replicates.

| [ATP] (mM) | $K_M$ (μM) | | $k_{cat}$ ($s^{-1}$) | | $k_{cat}/K_M \times 10^3$ ($M^{-1}s^{-1}$) | |
| --- | --- | --- | --- | --- | --- | --- |
| | hAMPD2-2 | hAMPD2-2Δ128 | hAMPD2-2 | hAMPD2-2Δ128 | hAMPD2-2 | hAMPD2-2Δ128 |
| 0 | N/D | N/D | N/D | N/D | N/D | N/D |
| 0.5 | $1.7 \times 10^3 \pm 200$ | $1.9 \times 10^3 \pm 300$ | $14.5 \pm 0.3$ | $190 \pm 5$ | $9 \pm 1$ | $100 \pm 20$ |
| 1 | $810 \pm 40$ | $770 \pm 50$ | $14.0 \pm 0.1$ | $180 \pm 2$ | $17 \pm 1$ | $230 \pm 20$ |
| 2 | $350 \pm 20$ | $460 \pm 20$ | $12.7 \pm 0.1$ | $170 \pm 1$ | $36 \pm 2$ | $390 \pm 20$ |
| 5 | $240 \pm 10$ | $210 \pm 10$ | $12.4 \pm 0.1$ | $150 \pm 1$ | $44 \pm 2$ | $750 \pm 30$ |

To determine the binding constant of ATP, different concentrations of ATP were added to 1 mM AMP-saturated hAMPD2-2. Curves were fit using a hyperbolic binding equation in GraphPad Prism, resulting in a calculated K_d_ = 1,113 ± 300 μM and 785 ± 200 μM (**Figure 3A** and **B**), for the full-length and catalytic domain, respectively. These values are lower than the average ATP concentration in liver,^20^ indicating that ATP is always bound to hAMPD2-2 under physiological conditions.

**Figure 3.**
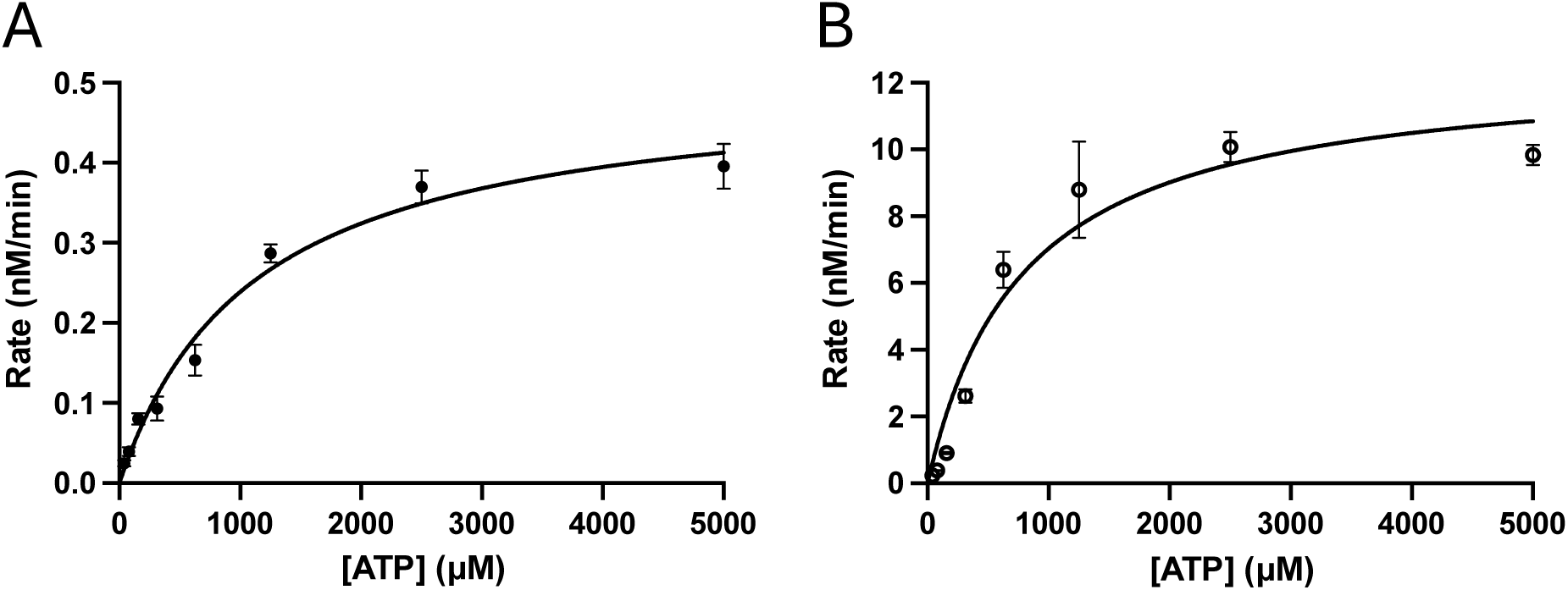
ATP binding curves for full-length and catalytic domain hAMPD2-2, at saturating AMP (1 mM). Rates of (**A**) Full-length and (B) catalytic domain at saturating [AMP]. Data points are from Figure S3 (at 0 mM P_i_) and fit to a hyperbolic binding equation.

### GTP inhibits hAMPD2-2 in the presence of ATP

GTP (at 2 mM) increases the T_m_ of both constructs by 7 °C compared to the unliganded enzyme (**Figure 2**). The effect is similar to that observed for ATP (2 mM) for both the full-length and catalytic domain of hAMPD2-2, which increases the T_m_ by 3 and 6 °C, respectively, compared to the no nucleotide sample. Both the full-length and the catalytic domain of hAMPD2-2 are inhibited by GTP in a concentration-dependent fashion (**Figures 4A** and **B**, **Table 2**). As discussed, in the absence of ATP, activity is negligible; therefore, the effect of GTP was assessed in the presence of saturating ATP (2 mM). A global model of inhibition was used to fit the data and to confirm the mode of inhibition as competitive (see Methods); for full-length *α* = 17, and for the catalytic domain *α* = 9. The results are consistent with GTP acting as a competitive inhibitor with AMP substrate, with K_i_ values of 74 ± 6 μM for the full-length and 101 ± 8 μM for the catalytic domain. The K_i_ values are the same for the full-length and catalytic domains, consistent with GTP binding at the active site in the catalytic domain. Inhibition of full-length hAMPD2-2 at 1 mM GTP was not plotted due to low signal-to-noise, where ΔA340 < 0.001, indicating the full-length enzyme is inactive at 1 mM GTP. A 10- to 100-fold difference in catalytic efficiency is observed between the two constructs (**Table 2**), with the catalytic domain exhibiting higher turnover and catalytic efficiency than the full-length enzyme at each GTP concentration. This further suggests an effect attributable to the N-terminal domain, as observed in the absence of GTP (**Table 1**).

**Figure 4.**
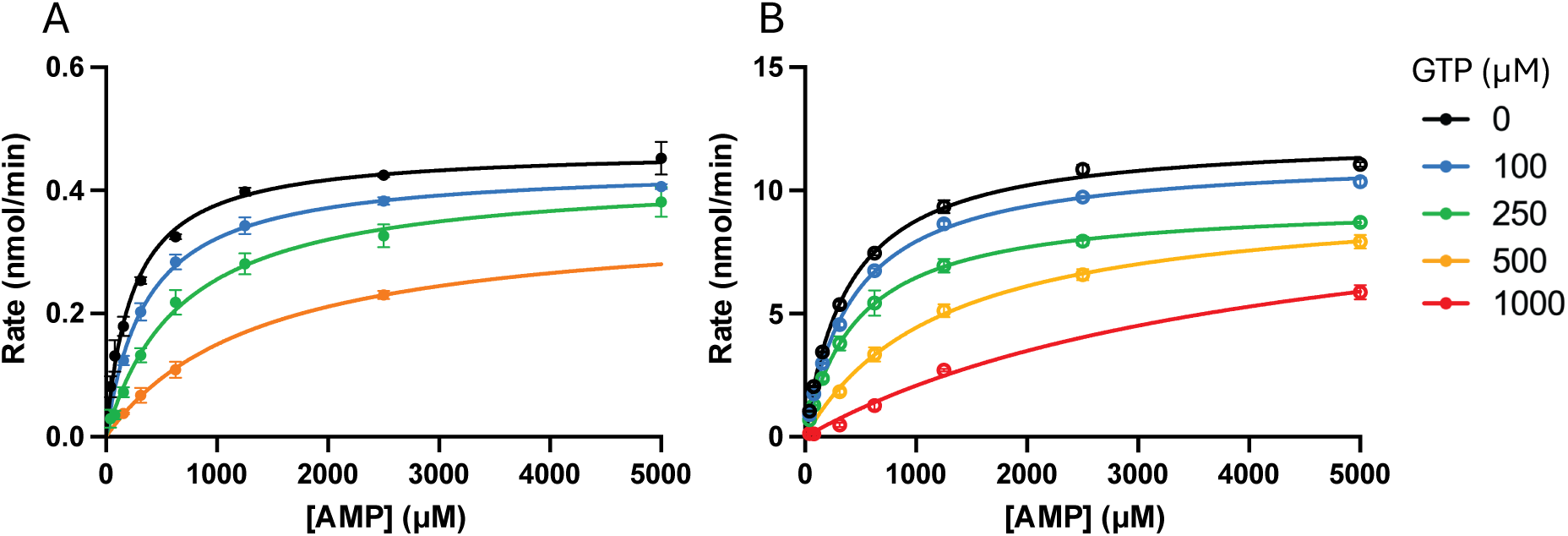
Competitive inhibition by GTP of ATP-activated hAMPD2-2 activity with substrate AMP. (A) Full-length hAMPD2-2 and (B) hAMPD2-2^Δ^_128_. [ATP] was constant at 2 mM. Steady-state kinetics data were fit using global fit in GraphPad Prism. Each data point was run in triplicate.

**Table 2.**
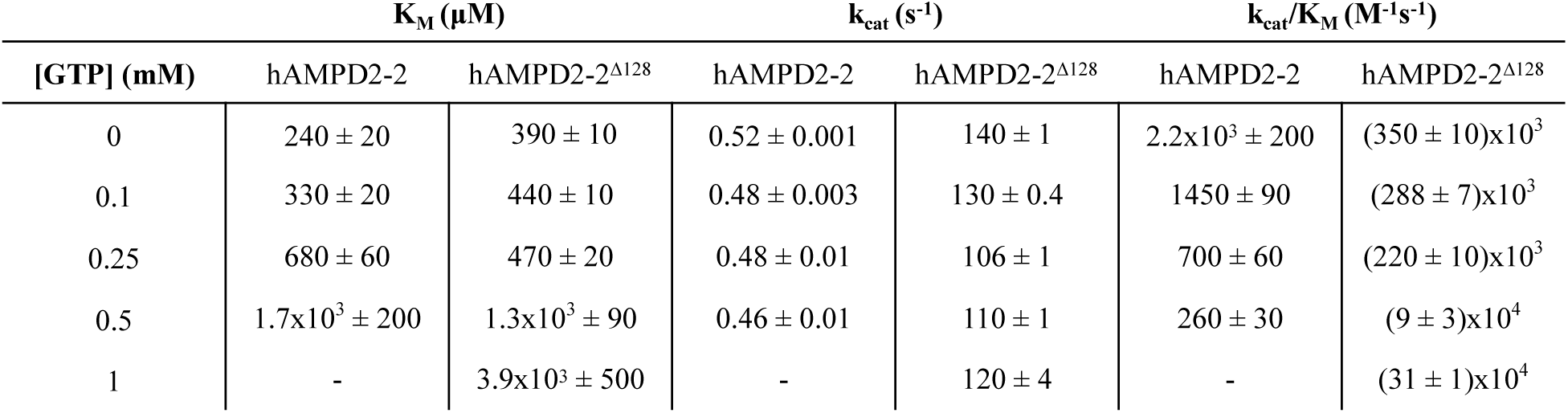
Steady-state kinetic constants of activated enzyme at varying [GTP]. ATP concentration is constant at 2 mM. Values are the mean determinations ± the standard deviation from 3 replicates.

### P_i_ inhibition of hAMPD2-2

Although we observed an effect of both ATP and GTP on the thermal stability of hAMPD2-2, P_i_ at 2 mM had no effect on thermal stability (T_m_ of 57 ± 1 °C, similar to that of the no nucleotide control of 56.1 ± 0.4 °C) (catalytic domain, data not shown). P_i_ has been shown to inhibit yeast AMPD at 0.35 mM in the presence of 0.1 mM ATP.^15^ However, ATP-activated hAMPD2-2 is not inhibited by P_i_ at the micromolar level (**Figure S3**). Instead, hAMPD2-2 shows inhibition by P_i_ at 10 mM (**Figure 5A**, **Table 3**). These results are similar for both the full-length enzyme and the catalytic domain, consistent with P_i_ binding at one or more sites within the catalytic domain. In addition, global fitting of the data indicates that both α values are > 1 (full length = 36; catalytic domain = 9), confirming competitive inhibition of Pi by the substrate AMP. The catalytic efficiency, k_cat_/K_m_ of the full length and the catalytic domain of hAMPD2-2 decrease 6- and 24-fold, respectively at high P_i_ concentrations (50 mM) (**Table S1**). This effect is reflected in K_m_ as expected for a competitive inhibitor. When fit to a competitive model with AMP substrate, the inhibition constant for P_i_ does not differ significantly between full-length (K_i_ = 10 ± 1 mM) and the catalytic domain (K_i_ = 3.7 ± 0.2 mM) of hAMPD2-2. The binding affinity is 10 to 30-fold lower than that previously observed in yeast AMPD.^15^ However, the studies on yeast AMPD were performed at 0.1 mM ATP, a concentration at which the activity was nearly undetectable in both the full-length and catalytic domains of hAMPD2-2. At [ATP] ∼ 0.1 mM we did observe a twofold decrease in activity comparing 0.5- and 5 mM P_i_ (vide infra, **Figure 10B**).

**Figure 5.**
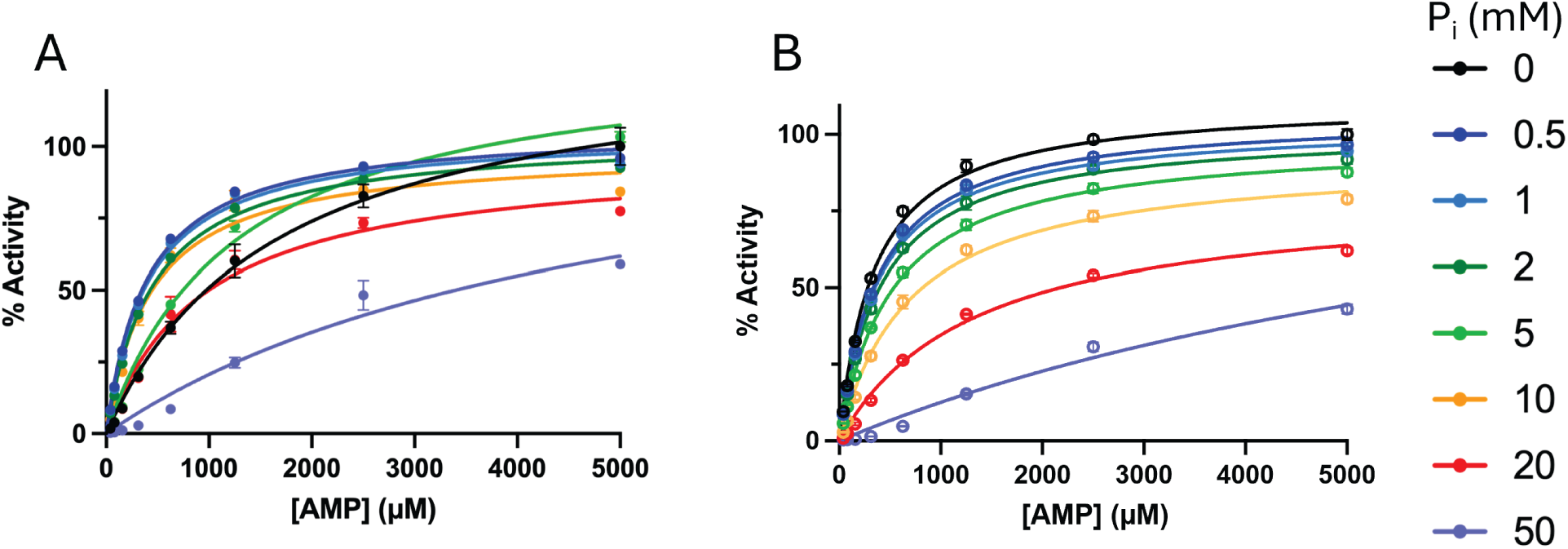
Competitive inhibition of P_i_ on activated hAMPD2-2 and hAMPD2-2^Δ128^. Panel A, full-length, panel B, catalytic domain. [P_i_] = 0 - 50 mM was used with activated enzyme (2 mM ATP).

**Table 3.** Steady-state kinetic constants of activated enzyme at varying [P_i_]. ATP concentration is 2 mM. Values are the mean determinations ± the standard deviation from 3 replicates.

| [$P_i$] (mM) | $K_M$ (μM) | | $k_{cat}$ ($s^{-1}$) | | $k_{cat}/K_M$ ($M^{-1}s^{-1}$) | |
| --- | --- | --- | --- | --- | --- | --- |
| | hAMPD2-2 | hAMPD2-2Δ128 | hAMPD2-2 | hAMPD2-2Δ128 | hAMPD2-2 | hAMPD2-2Δ128 |
| 0 | 350 ± 20 | 340 ± 20 | 12.3 ± 0.1 | 140 ± 1 | (36 ± 12)$\times 10^3$ | (41 ± 2)$\times 10^4$ |
| 0.5 | 390 ± 10 | 380 ± 10 | 12.3 ± 0.1 | 135 ± 1 | (315 ± 8)$\times 10^2$ | (360 ± 10)$\times 10^3$ |
| 1 | 400 ± 20 | 380 ± 10 | 12.2 ± 0.1 | 132 ± 1 | (30 ± 2)$\times 10^3$ | (350 ± 10)$\times 10^3$ |
| 2 | 460 ± 20 | 410 ± 10 | 12.0 ± 0.1 | 129 ± 1 | (26 ± 1)$\times 10^3$ | (313 ± 8)$\times 10^3$ |
| 5 | 1300 ± 100 | 520 ± 20 | 39.4 ± 0.6 | 125 ± 4 | (31 ± 2)$\times 10^3$ | (24 ± 1)$\times 10^4$ |
| 10 | 430 ± 40 | 700 ± 40 | 28.9 ± 0.4 | 117 ± 1 | (67 ± 2)$\times 10^3$ | (170 ± 10)$\times 10^3$ |
| 20 | 900 ± 100 | 1300 ± 100 | 28.5 ± 0.5 | 119 ± 4 | (30 ± 4)$\times 10^3$ | (31 ± 8)$\times 10^3$ |
| 50 | 5000 ± 1000 | 9000 ± 2000 | 36 ± 2 | 119 ± 4 | (7 ± 1)$\times 10^3$ | (31 ± 3)$\times 10^3$ |

To determine the inhibition mechanism of P_i_ on ATP-activated hAMPD2-2, assays were performed using saturating AMP (1 mM) and varying the concentration of ATP and P_i_ (**Figure S3**). Results were fit to a four-parameter Hill plot binding curve and normalized to enzyme activity at 5 mM ATP at each P_i_ concentration. For full-length enzyme, as P_i_ increases (from 0 to 5 mM), the concentration of ATP needed to reach 50% activity (EC_50_ value) increases 1.5-fold. The effect on ATP affinity is also small for hAMPD2-2^Δ^_128_. These results are consistent with weak binding of P_i_ to hAMPD2-2. Addition of 50 mM P_i_ (25 times the physiological concentration in liver cells^20^) shows a greater shift in apparent ATP affinity (higher EC_50_ value) in full-length enzyme. More specifically, enzymatic activity is observed only at > 1.25 mM ATP, suggesting that Pi competes with ATP and therefore acts as an allosteric inhibitor at low ATP concentrations.

Further investigation of P_i_ inhibition of ATP-activated hAMPD2-2 was conducted using increasing amounts of Pi at lower, but saturating, concentrations of AMP and ATP (1 mM each) (**Figure 6**). Fraction enzyme inhibition *i* was measured and fit to a competitive inhibition model with AMP to determine K_i_ values for full-length enzyme using Equation 5 (see Methods), K_i_ = 1.9 ± 0.4 mM and catalytic domain, K_i_ = 5 ± 1 mM. These values are similar to those determined using 2 mM ATP (**Figure 5**), with Ki values of 10 mM and 3.7 mM for the full-length and catalytic domains, respectively. These affinities are equal to or somewhat above the physiological concentrations of P_i_.^21^

**Figure 6.**
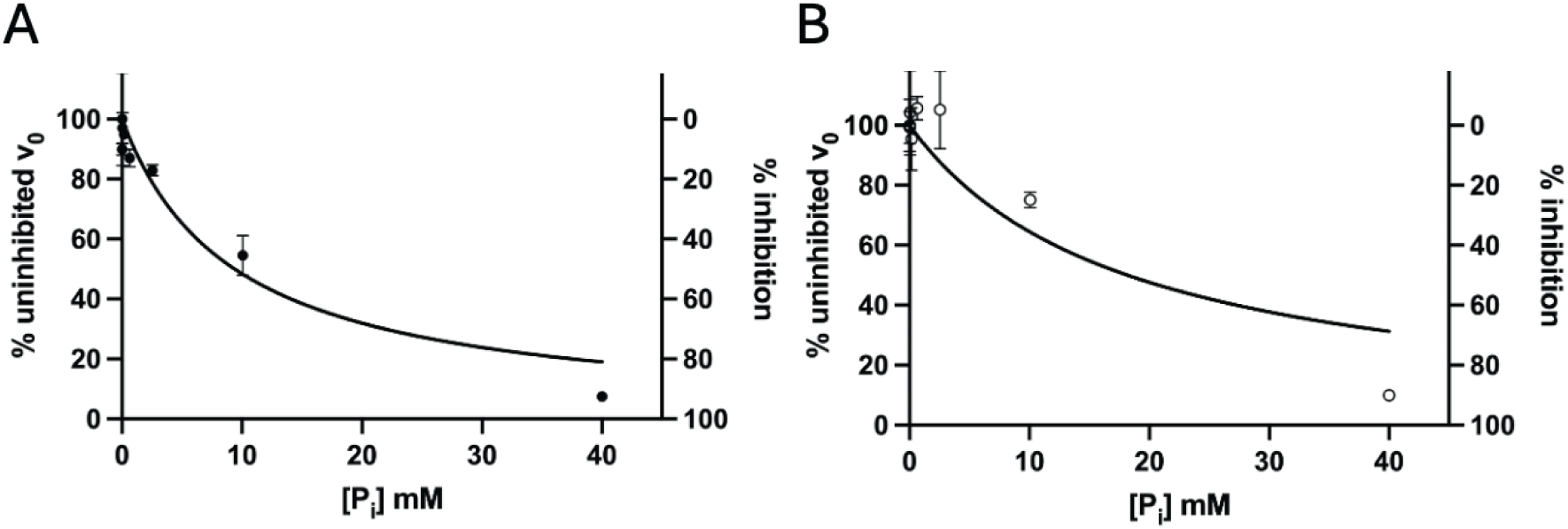
Activated hAMPD2-2 is inhibited at high P_i_ concentrations. Enzyme activity was measured in the presence of 1 mM AMP and 1 mM ATP and results fit to Equation 5 (assuming competitive inhibition with AMP, see Methods).

### The role of P_i_ in hAMPD2-2 inhibition in the presence of GTP

Further work investigated the action of P_i_ in the presence of GTP. Studies by Van de Berghe and colleagues^16^ on mouse liver AMPD and by Merkler & Schramm^17^ on yeast AMPD suggest P_i_ increases GTP inhibition when present. This could conceivably be achieved by allosteric or orthosteric binding. To assess the model, assays were performed using varying amounts of GTP, with saturating AMP (1 mM) and ATP (2.5 mM), in the absence and presence of P_i_. AMPD2-2 is fully activated at 2.5 mM ATP, which is within the physiological concentration range in liver cells.^20^ Including 2 mM P_i_ inhibits hAMPD2-2 to a greater extent than GTP alone (**Figure 7 and S4**), by ∼ two to three-fold. Results for the full-length and catalytic domain are similar. Previous reports examining the effects of ATP, GTP and P_i_ confirm an increase in inhibition in mouse liver AMPD when both GTP and P_i_ are present at physiological concentrations.^16^ However, the mechanisms of the effects of these regulators on AMPD remained unclear. To assess the mode of inhibition of hAMPD2-2, the effect of varying P_i_ on activity in the presence of GTP was tested under AMP-saturated, ATP-activated conditions (**Figure S5**). We observe a 10-fold decrease in enzyme activity as Pi concentration increases for both hAMPD2-2 constructs (**Figure 8**).

**Figure 7.**
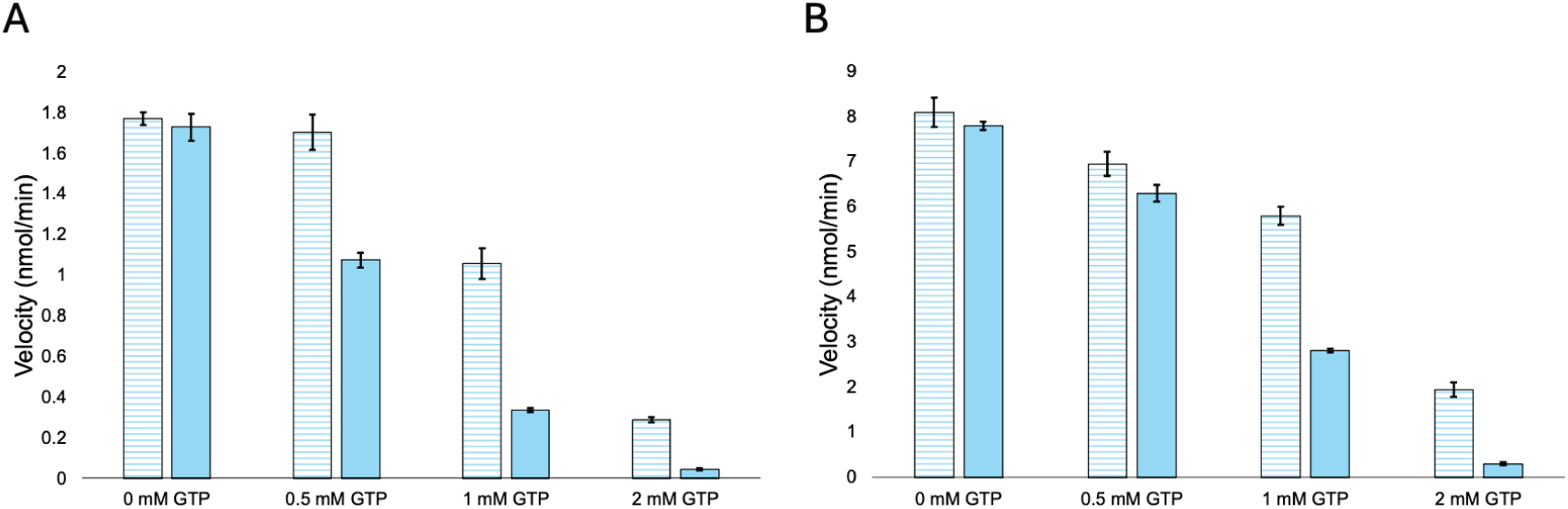
Effect of P_i_ on hAMPD2-2 and catalytic domain activity when inhibited by GTP. Panel A, full-length, panel B, catalytic domain. Horizontal stripped bars and filled bars correspond to 0 mM P_i_ and 2 mM P_i_, respectively. Data points were obtained from Figure S4 at 2.5 mM ATP.

**Figure 8.**
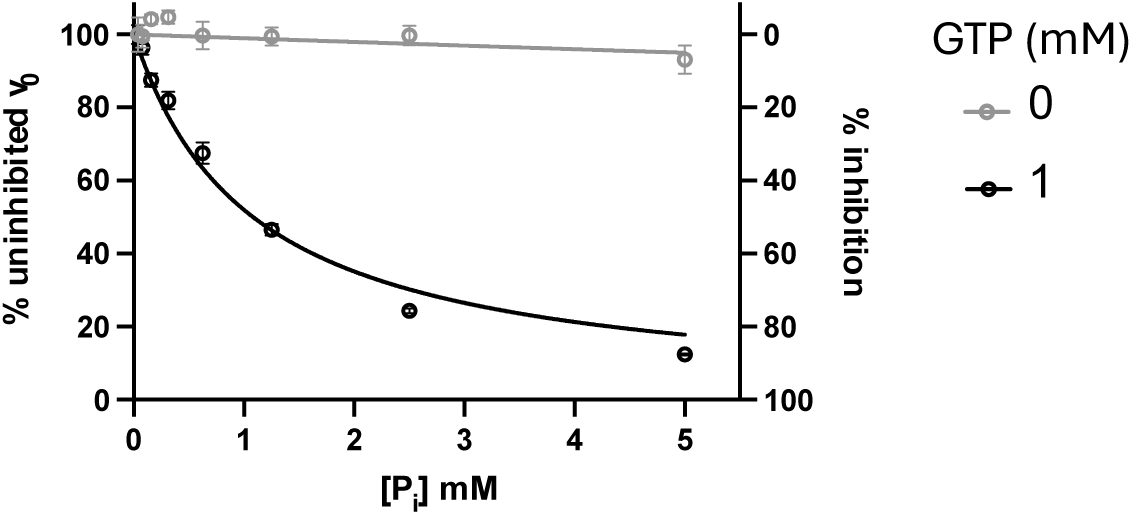
Effect of GTP on saturated and activated catalytic domain of hAMPD2-2, at increasing levels of P_i_. Data points were fitted to Equation 5 (see Methods), curves were fitted assuming competitive inhibition with AMP. Data was determined at 1 mM AMP and 2 mM ATP. Absence of GTP is shown in grey circles, presence of GTP (1 mM) is shown in black circles.

Because the high concentrations of ATP in this experiment may obscure effects resulting from P_i_ binding to the allosteric site, the dose-response curve was repeated with 1 mM ATP (**Figure 6**). As there is virtually no activity in the absence of ATP, competing with ATP is indistinguishable from competing with substrate AMP, and the P_i_-binding data could be fit to a competitive inhibition model. The K_i_ value for P_i_, measured in the presence of GTP was 1.3 ± 0.1 mM, which is five-fold lower than the K_i_ in the absence of GTP (5 mM, **Figure 9**). Taken together, the data are consistent with a model in which P_i_ can bind to either the active site or allosteric site, and where occupancy of the allosteric site enhances ligand binding to the active site and vice versa.

## DISCUSSION

Previous studies^12^ have characterized isozymes of the liver isoform of human AMPD by expression and purification from Sf9 cells. Although full-length yeast AMPD has been purified heterologously from *E. coli*, rapid loss of enzyme activity and auto-proteolysis of the N-terminal domain have prevented further studies of the native construct. Herein, we report the successful expression and purification of both the full-length and catalytic domains of the hAMPD2-2 isozyme from *E. coli* (**Figure S6**). Throughout our studies, the catalytic activity of the full-length enzyme was assessed to ensure the integrity of the protein.

Because of the instability of the full-length construct, studies in yeast AMPD were mainly performed on the catalytic domain,^15, 17, 22–24^. These findings for yeast full-length AMPD show 100-fold lower specific activity compared to the proteolyzed enzyme, consistent with our demonstration of higher specific activity for the truncated construct of hAMPD2-2.^15^

In the cell, under normal conditions, ATP is present in millimolar concentrations (in liver, 2-5 mM).^20^ Our results show that in the absence of either GTP or P_i_, hAMPD2-2 is fully activated at these physiological ATP concentrations, for both full-length and truncated hAMPD2-2. In addition, the presence of the N-terminal domain does not affect the activation of the enzyme by ATP. However, it does decrease the enzyme turnover rate, suggestive of either a conformational change in this domain that prevents access of the AMP substrate to the active site or prevents activator ATP binding.

Previous studies of the catalytic domain of yeast AMPD show that GTP is an inhibitor, competitive with the AMP substrate of AMPD.^15^ Here, we confirm these results for both the full-length hAMPD2-2 and the catalytic domain of the human enzyme with a K_i_ for GTP of 74 and 101 μM, respectively (**Figure 4**). These values are lower than the average GTP concentration in cells of 500 μM,^20^ supporting a model where GTP inhibition of AMPD2 occurs in liver cells under physiological conditions. However, during fructose consumption, the amount of available GTP decreases in the cell as it is consumed by nucleoside diphosphate kinase (NDPK), which maintains ATP homeostasis.^25–27^ This metabolic change would relieve the inhibition of hAMPD2-2 by GTP, thereby increasing uric acid.^28^

When ATP is present at normal physiological concentrations (2 mM in liver),^20^ the effect of P_i_ *alone* on AMPD2 would be negligible. Reduction in AMPD2 activity would only be observed when P_i_ concentrations reached ≥ 10 mM (**Figure 5**); this inhibitory effect has previously been observed in trout gill AMPD (79% sequence identity with hAMPD2-2).^29^ In our work, the activation of the enzyme by ATP is consistent (EC_50_ increases by less than two fold), regardless of the amount of P_i_ present (**Figure S3**). Indeed, there are no significant differences in catalytic efficiency when Pi is present at concentrations up to 5 mM (**Table S1**). Assuming competitive inhibition with AMP in the active site, the K_i_ values for P_i_ are 10 and 3.7 mM for the full-length and truncated hAMPD2-2 (**Figures 5A** and **5B**).

Physiological concentrations of P_i_ (2 mM) only have an observable effect in the presence of GTP (**Figures 8 and 9**). When both GTP and P_i_ are present, both inhibit hAMPD2-2; GTP competitively inhibits the substrate AMP, and Pi competes with the activator ATP. This model has not been previously proposed for AMPD. The effect of P_i_ in the presence of GTP is consistent with a model in which P_i_ is binding in the ATP site at physiological concentrations of P_i_ (**Figure S3**). This was confirmed by fitting data from **Figure S3B** (catalytic domain) to an inhibition model incorporating competitive inhibition with the enzyme activator ATP (**Figure 10**). Our data confirmed competitive inhibition of GTP with AMP, but the possibility should be considered where the purine GTP could also bind to the ATP activation site, as was seen for P_i_. Therefore, data for GTP inhibition of activated hAMPD2-2 (**Figure S4A** and **S4C**) was also fit to Equation 6; the lack of fit to this kinetic model confirms GTP does not bind to the activation site.

Global fitting in the program KinTek was performed to determine whether traces from the steady-state experiments could be described within a single kinetic framework (**Figures S7, S8** and **S9**), resulting in a comprehensive mechanism for hAMPD2-2 (**Figure 11**). The resulting microscopic rate constants determined for the catalytic domain of hAMPD2-2 were broadly consistent with the steady-state analysis: K ^AMP^ = 471.4 ± 11.9 µM, k_cat_ = 133.0 ± 1.7 s⁻¹ and K_D_ ^GTP^ = 60.8 ± 12.6 µM. These values are similar to those determined using steady state fit, where K ^AMP^ = 460 ± 20 µM, k_cat_ = 170 ± 1 s⁻¹ and K_i_ ^GTP^ = 101 ± 8 µM. Because the available data were insufficient to uniquely constrain all microscopic rate constants, the global fit was not interpreted mechanistically but was instead used as an internal consistency check of the overall kinetic behavior.

**Figure 10.**
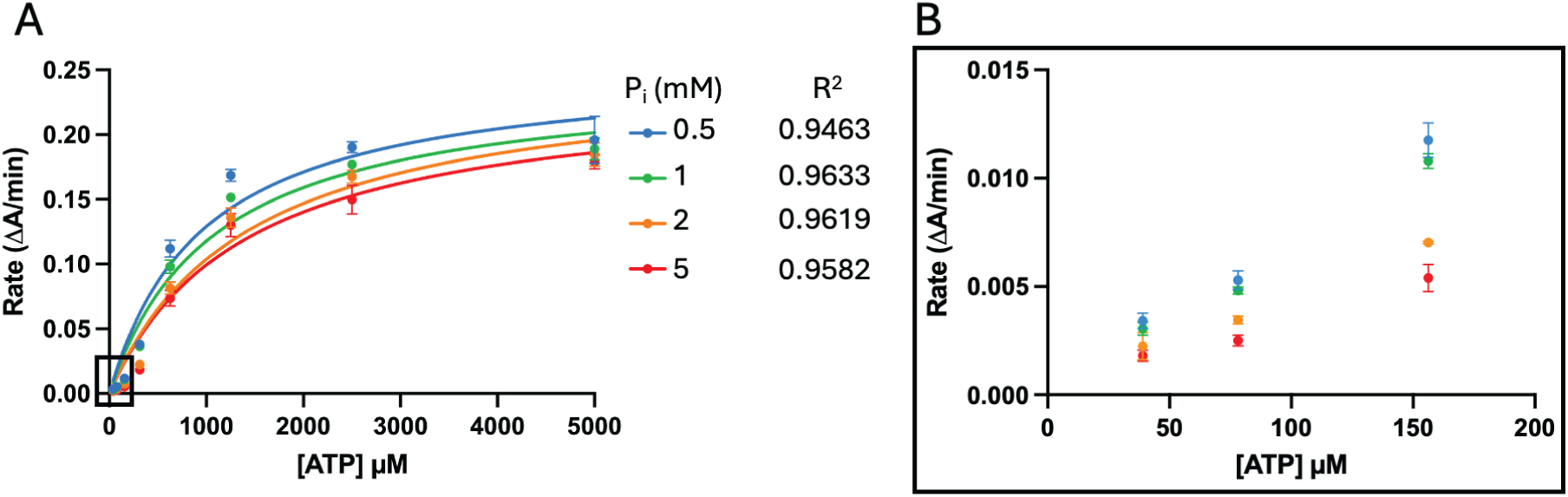
P_i_ competes with ATP under AMP saturated conditions, inhibiting hAMPD2-2^Δ128^. Curves were fit to an inhibition model (see Methods, Equation 6) suggesting P_i_ inhibits hAMPD2-2 *via* competitive inhibition of ATP at [AMP] = 1.25 mM (from Figure S4B). Panel A, [ATP] = 39 μM - 5 mM. Panel B, scale-up view of [ATP] range from 39 μM - 156 μM.

**Figure 11.**
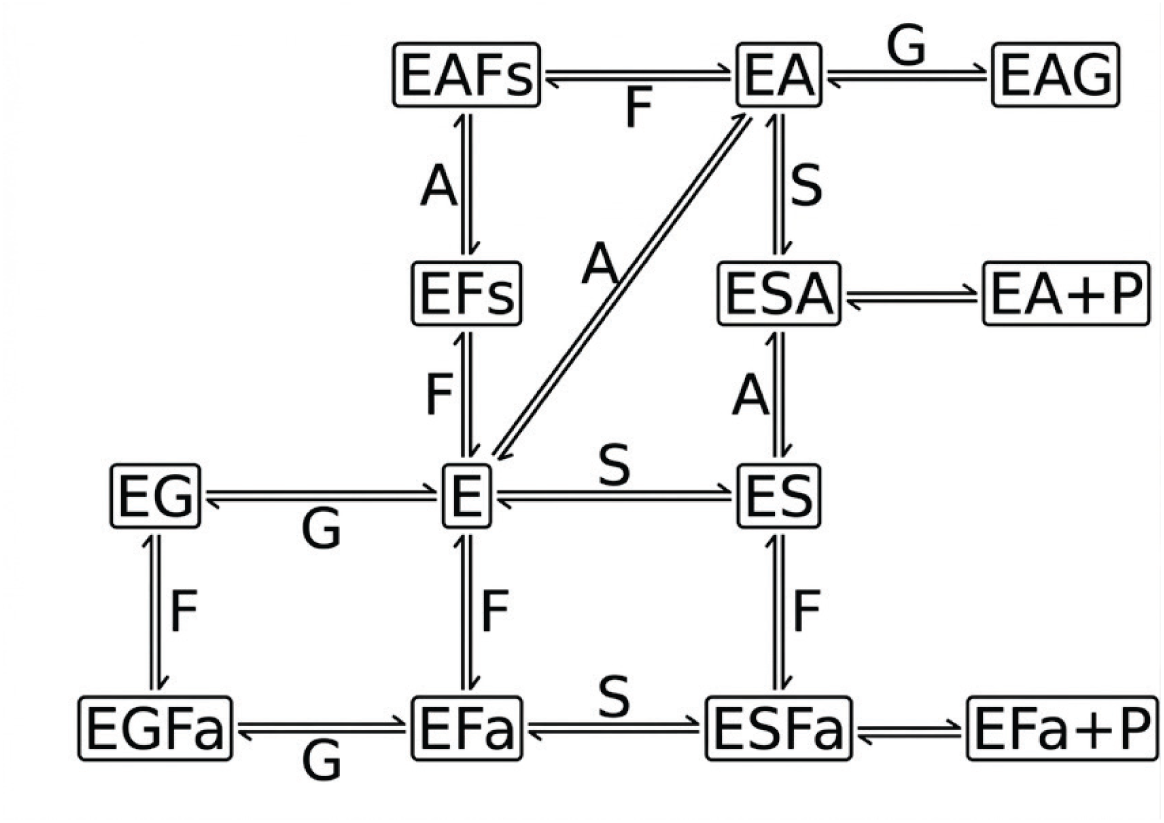
Proposed mechanism of hAMPD2-2 used in global fitting. E, enzyme; S, AMP; A, ATP; G, GTP; Fs, P_i_ bound at the catalytic site; Fa, P_i_ bound at the allosteric site; P, product. The scheme was drawn using Google Gemini.

Studies by Van den Berghe *et al*. (1977) on the rat liver AMPD orthologue (rAMPD2, 97.9% sequence identity) were the first to show how all three regulators affect the enzyme, with ATP as an activator and GTP and P_i_ both inhibiting AMPD in tandem,^16^ but it is unclear if the rAMPD2 used is full-length or if the N-terminal domain auto-proteolyzed. Here, we confirm the mechanism of all three effectors of hAMPD2-2 under physiological conditions, and under conditions mimicking that of fructose consumption. Our findings are consistent with a model in which the allosteric site is not located in the N-terminal domain. All results were similar between full-length and the catalytic domain construct, other than the higher catalytic efficiency and turnover for the truncated enzyme. Previous studies of yeast AMPD2 show that ATP increases the affinity for AMP, decreasing the K_M_, in both full-length and catalytic-domain proteins.^17^ Here we confirm ATP binds to an allosteric site on the catalytic domain in human AMPD, as the presence of the N-terminal domain does not affect our results. The N-terminal does not play a regulatory role by binding to the tested metabolites.

We propose a model for pathway regulation in liver cells, based on the mechanism used in the global fit (**Figure 11**) for activity under normal conditions and during fructose consumption (**Figure 12**). Under normal physiological conditions, hAMPD2-2 is inhibited by GTP and P_i_. With fructose consumption, ATP, GTP and P_i_ concentrations decrease;^3, 30^ the general P_i_ pool decreases due to the action of hKHK-C to phosphorylate fructose *via* ATP at the start of the pathway.^7^ This decrease in cellular ATP forces NDPK to utilize other nucleotide triphosphates, such as GTP, for phosphorylating ADP into ATP in order to maintain ATP homeostasis.^26, 27^ Concurrently, GDP is not phosphorylated into GTP, leading to GTP concentrations decreasing in the cell. With less P_i_ and GTP available to inhibit hAMPD2-2, the catalytic efficiency of the enzyme increases 10-fold, thereby increasing IMP and uric acid levels (**Figure 13**).

**Figure 12.**
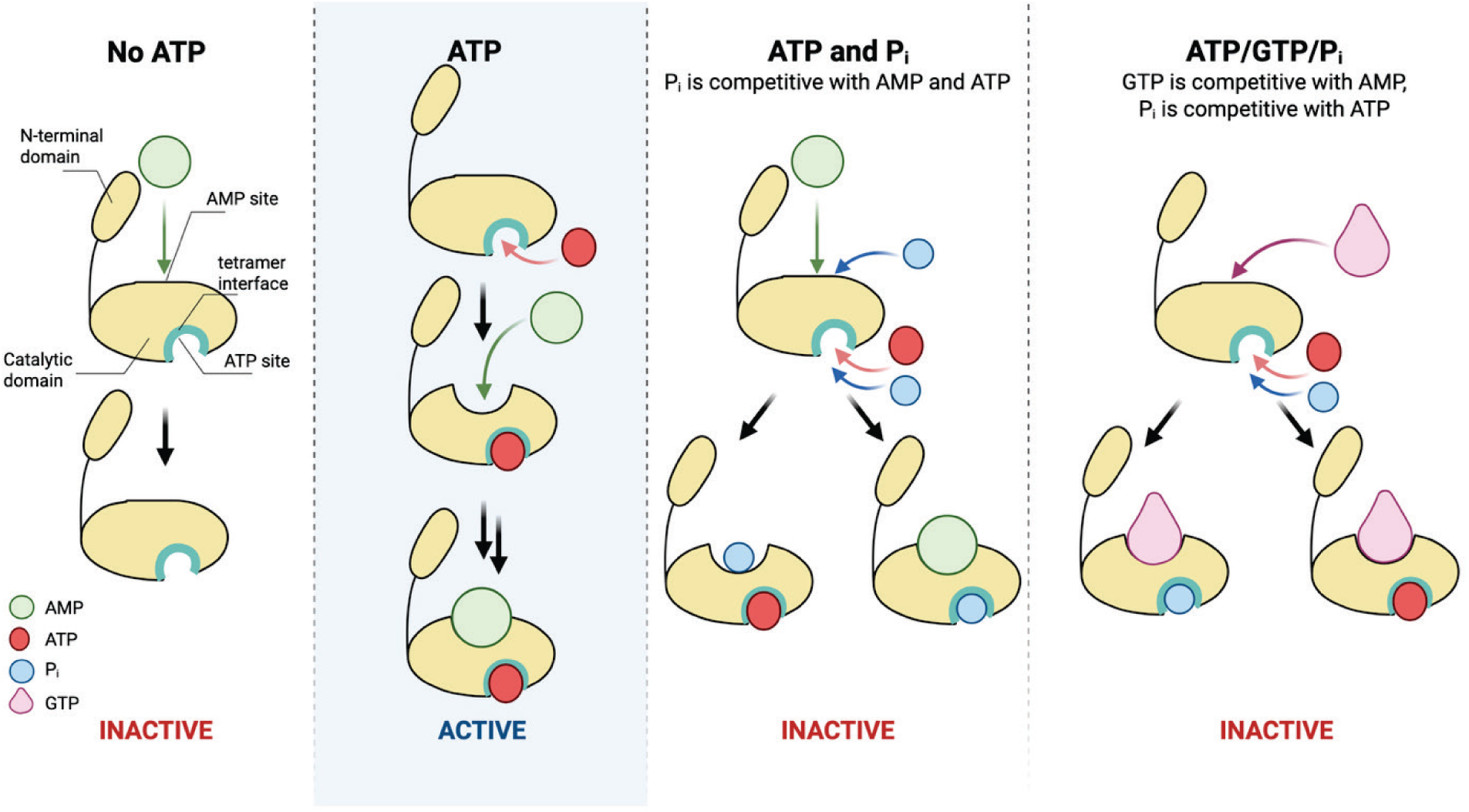
Activation and inhibition model of monomeric hAMPD2-2. ATP-activation is necessary for hAMPD2-2 catalysis. P_i_ competes with both AMP and ATP binding sites, inhibiting hAMPD2-2. GTP competes with the AMP site, whereas P_i_ competes with the ATP binding site, enhancing enzyme inhibition. The enzyme is a tetramer; a single protomer is diagrammed.

**Figure 13.**
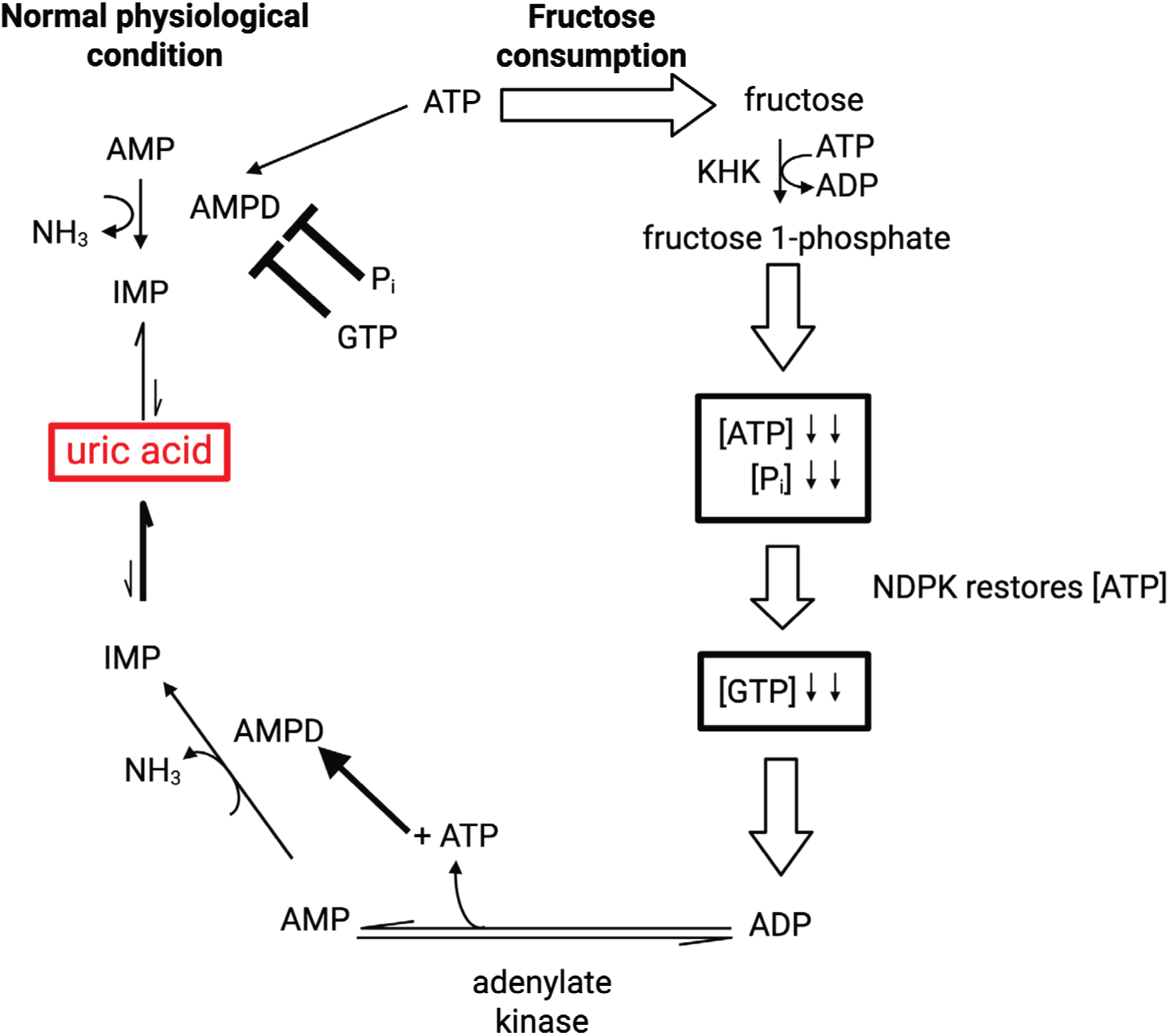
hAMPD2-2 activation during fructose consumption. Under normal physiological conditions, liver AMPD is inhibited by GTP and P_i_, limiting the concentration of cellular uric acid. With fructose consumption, ATP and GTP concentrations decrease concomitantly, leading to increased nucleoside diphosphate kinase activity, thereby activating AMPD and increasing uric acid.

## CONCLUSIONS

AMPD-2 acting in the liver and brain serves as a critical metabolic "rheostat" at the intersection of energy sensing, purine metabolism, and fat accumulation.^31^ Hence, the influences of its allosteric effectors, ATP, GTP, and P*_i_* are complex. This study elucidates the interplay among these effectors in the human enzyme and supports a model for how this important enzyme is regulated. Our results are consistent with the action of effectors from previous work on AMPD3 and the yeast enzyme.^32, 33^ An important difference was that throughout the investigation, both the full-length enzyme and the more widely studied truncated version were included, making it clear that the binding site for these effectors is not on this N-terminal domain, whose role remains unclear. Given the clear requirement for ATP for activity, and the homeostatic levels of ATP in the cell, the dampening effect of this N-terminal domain on overall activity may be important for limiting purine degradation when not required. Additionally, we describe the interaction of AMPD and its activators and inhibitors, quantitatively positing a dual role for P*_i_* and a more important role for GTP than previously appreciated.

A deeper understanding of AMPD2 is essential, particularly for the development of allosteric inhibitors.^34^ Excessive fructose intake rapidly depletes hepatic ATP, activating AMPD2 and driving the overproduction of uric acid. This cascade contributes to mitochondrial dysfunction and suppresses AMP-activated protein kinase (AMPK)—the body’s central metabolic regulator that promotes fat oxidation and limits fat synthesis—ultimately predisposing individuals to Metabolic Dysfunction-Associated Steatotic Liver Disease.^31, 35^ An effective allosteric inhibitor could, in principle, “trick” the liver into a fasted, fat-burning metabolic state even under high-calorie conditions.

In the brain, such inhibitors may influence feeding behavior^36^ and help preserve AMP levels, potentially offering protection against neuroinflammation and providing leads for research into neurodegenerative diseases.^37^ Targeting AMPD2 therefore represents a promising strategy for counteracting the cellular consequences of high-sugar, high-fat diets and may ultimately contribute to therapies for widespread metabolic disorders, including MASLD, obesity, dementia, and type 2 diabetes.

## AUTHOR INFORMATION

### Corresponding Authors

Karen N. Allen – Department of Chemistry, Boston University, Boston, MA 02215, United States; orcid.org/0000-0001-7296-0551;

Dean R. Tolan – Department of Biology, Boston University, Boston, MA 02215, United States; orcid.org/0000-0002-0598-7241;

### Authors

Ashley M. Rebelo – Department of Chemistry, Boston University, Boston, MA 02215, United States; orcid.org/0000-0001-6691-6124

Nemanja Vuksanovic – Department of Chemistry, Boston University, Boston, MA 02215, United States; orcid.org/0000-0003-2883-6312

Lanlan Han – Department of Chemistry, Boston University, Boston, MA 02215, United States; orcid.org/0009-0007-8050-1882

### Present Addresses

Ashley M. Rebelo – Department of Biology, Massachusetts Institute of technology, Cambridge, MA 02139, United States;

Lanlan Han – Department of Biosciences and Bioinformatics, Xi’an Jiaotong-Liverpool University, Suzhou 215123, China;

Nemanja Vuksanovic – Department of Chemistry, Boston University, Boston, MA 02215, United States;

### Author Contributions

A.M.R. performed expression, purification, steady-state kinetics and data analysis of hAMPD2 and wrote the manuscript. L.H. identified initial expression conditions for His_6_-SUMO tagged hAMPD2-2^Δ^_128_, initial purification conditions of this construct and designed the preliminary coupled enzyme assay. N.V. performed Kintek fitting to obtain the global fit of the steady-state parameters. K.N.A. and D.R.T. supervised the study and wrote and edited the manuscript.

### Funding Sources

Supported by NIH grant U01AA027997 (to DRT and KNA).

### Notes

DRT has equity with Colorado Research Partners LLC, which is developing KHK inhibitors. AMR, NV, LLH, and KNA declare no competing financial interest.

## Supporting information

Supplement containing Supporting methods, Figs S1-S9, Tables S1-S3

## ACKNOWLEDGMENTS

We thank Dr. Lizbeth Hedstrom from the Department of Chemistry at Brandeis University for gifting the cpIMPDH coupling enzyme construct plasmid used for the kinetic studies.

## ABBREVIATIONS

AMD,: yeast adenosine 5’-monophosphate deaminase;
cpIMPDH,: inosine 5’-monophosphate dehydrogenase from *Cryptosporidium parvum*;
Fru 1P,: fructose 1-phosphate;
hAMPD,: human adenosine 5’-monophosphate deaminase;
HFI,: hereditary fructose intolerance;
hKHK,: human ketohexokinase;
HEPPS,: 3-[4-(2-Hydroxyethyl)piperazin-1-yl]propane-1-sulfonic acid;
IMP,: inosine 5’-monophosphate;
TEA,: triethanolamine.

