## Supplement containing Supporting methods, Figs S1-S9, Tables S1-S3 for "Deciphering AMP deaminase-2 structure, activators and regulators underpinning cellular function in human fructose and nucleotide metabolism"

**Table of contents**

**Supplementary Methods**
- cpIMPDH expression and purification
- cpIMPDH activity assay

**Supplementary Figures**
- S1. Sequence alignment of full length hAMPD isoforms and isozymes
- S2. ATP activates hAMPD2-2 and hAMPD2-2$^{\Delta128}$
- S3. Addition of $P_i$ with AMP-saturated full-length (Panel A) and catalytic domain of hAMPD2-2 (Panel B) under varying [ATP]
- S4. Hill plots of substrate saturated hAMPD2-2 in the absence and presence of 2 mM $P_i$ with increasing concentrations of ATP, varying [GTP]
- S5. Effect of GTP on saturated and activated hAMPD2-2, at increasing levels of $P_i$
- S6. Coomassie stained 12% SDS-Page of full-length and catalytic domain constructs of hAMPD2-2 purification from *E. coli*
- S7. Reduced global fitting of A340 traces across GTP, ATP and $P_i$ concentrations
- S8. Reduced global fits of AMP titrations under varying ATP and GTP conditions
- S9. Global fitting of ATP titrations at elevated $P_i$ concentrations

**Supplementary Tables**
- Table S1. Steady-state kinetic constants for activated enzyme with increasing $P_i$ concentrations
- Table S2. Titration of GTP with AMP saturated and ATP activated hAMPD2-2 in the absence and presence of $P_i$
- Table S3. Kintek global fitting model values

**Supplementary methods**

*cpIMPDH expression and purification*

pET28a including the gene encoding poly-histidine tagged IMP dehydrogenase from *Cryptosporidium parvum* (cpIMPDH) was generously donated by Professor Lizbeth Hedstrom, Brandeis University. This construct was modified from the original cpIMPDH sequence by deleting the 90-134 loop and replacing it with SGG to increase protein stability.[1] The plasmid was transformed into competent BL21(DE3) *E. coli* cells (New England Biolabs) and plated onto a Luria Broth-agar plate containing 50 µg/mL kanamycin. A starter culture of 250 mL Luria Broth was seeded with a single colony and grown overnight at 37 °C. Large scale 1-L cultures containing 50 µg/mL kanamycin were inoculated using a 1:50 dilution of the starter culture and grown at 37 °C until $OD_{600}$ reaches 0.7, induced with 0.25 mM IPTG and grown 14-16 hours at 25 °C. Cells were harvested at 5,383 x g for 20 min at 4 °C.

All purification steps were performed cold (4 °C). Frozen cells from either a 4 or 6-L growth were thawed and resuspended in lysis buffer (25 mM HEPPS pH 8, 500 mM KCl, 25 mM imidazole, 1 µM glutathione, EDTA-free protease inhibitor tablet (ThermoFisher) and 6 mg/100 mL resuspension of DNase I (GoldBio)). Cells were lysed *via* microfluidizer and clarified *via* ultracentrifugation at 106,255 x g for 35 minutes. The soluble fraction was purified using affinity chromatography *via* HisTrap High Performance Ni-Sepharose column (Cytiva), which was previously equilibrated in wash buffer (25 mM HEPPS pH 8, 500 mM KCl, 25 mM imidazole). After application of the soluble fraction, the column was washed with 5 column volumes (CV) of wash buffer. The protein was eluted from the column by a 2-step gradient using 50% and 100% elution buffer (25 mM HEPPS pH 8, 500 mM KCl, 275 mM imidazole). The eluted protein was concentrated, and buffer exchanged into storage buffer (25 mM HEPPS pH 8, 500 mM KCl, and

10% (v/v) glycerol). Protein was quantified *via* Bradford assay according to the manufacturer's instructions (ThermoFisher). Protein purity was assessed using SDS-PAGE. Aliquots were flash-frozen and stored at -80 °C.

*cpIMPDH activity assay*

The activity of cpIMPDH was determined using the absorbance of NADH at $\lambda_{340}$ in a colorimetric assay. Purified cpIMPDH (1, 5, and 10 µL at 10-25 mg/mL) was added to a cocktail containing 50 mM TEA pH 7.4, 4 mM $MgCl_2$, 2 mM ATP, 5 mM $NAD^+$, 1 mM DTT and 1 mM IMP (buffer based on kinetics assays done on yeast AMD[2,3]). Each concentration was run in triplicate in a Corning® 96 well plates (9018BC) in a final volume of 200 µL and at ambient temperature (between 24 – 26 °C) for 10 minutes and readings recorded every 10 seconds using a SpectraMax M5® plate reader (Molecular Devices). The $\Delta A_{340}$/time (min) was determined using the linear part of the collected data using SoftMax Pro software.

### Supplementary Figures

A

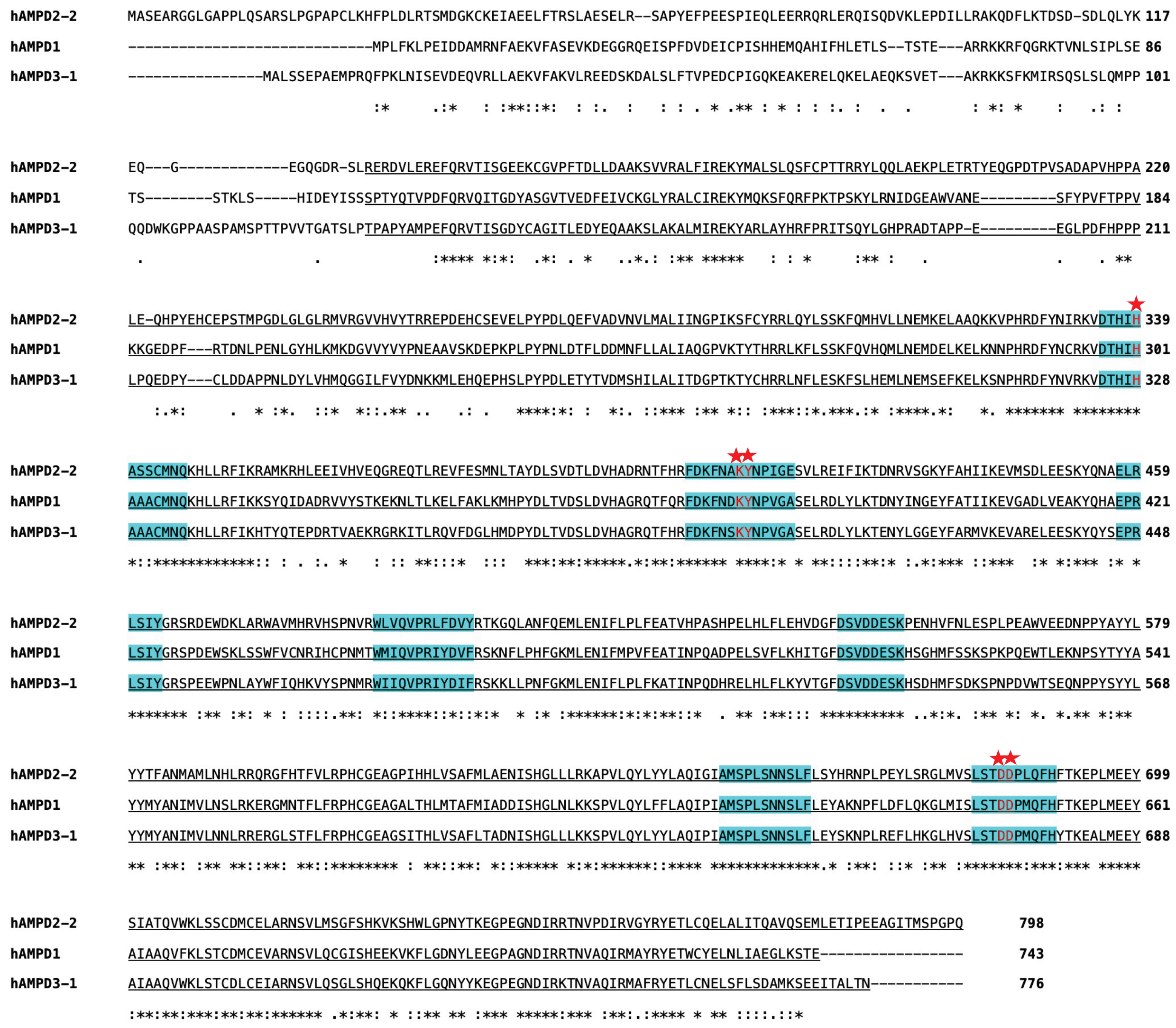

B

```
hAMPD2-2    --MASEARGGLGAPPLQSARSLPGPAPCL----KHFPLDLRTSMDGKCKEIAEELFTRSLAESELRSAPYEFPEESPIEQLEERRQRLERQISQDVKLEPDILLRAKQDFLKTDSDSDLQ 114
hAMPD2-3    MWQSQAPAGAAQTPPLSPPWSQPWHPIHLALASPRPNIPLRSGPACRPPLQLQELFTRSLAESELRSAPYEFPEESPIEQLEERRQRLERQISQDVKLEPDILLRAKQDFLKTDSDSDLQ 120
hAMPD2-5    -------------------------------------------MDGKCKEIAEELFTRSLAESELRSAPYEFPEESPIEQLEERRQRLERQISQDVKLEPDILLRAKQDFLKTDSDSDLQ 77
                                                          :     :*******************************************************************

hAMPD2-2    LYKEQGEGQGDRSLRERDVLEREFQRVTISGEEKCGVPFTDLLDAAKSVVRALFIREKYMALSLQSFCPTTRRYLQQLAEKPLETRTYEQGPDTPVSADAPVHPPALEQHPYEHCEPSTM 234
hAMPD2-4    LYKEQGEGQGDRSLRERDVLEREFQRVTISGEEKCGVPFTDLLDAAKSVVRALFIREKYMALSLQSFCPTTRRYLQQLAEKPLETRTYEQGPDTPVSADAPVHPPALEQHPYEHCEPSTM 240
hAMPD2-5    LYKEQGEGQGDRSLRERDVLEREFQRVTISGEEKCGVPFTDLLDAAKSVVRALFIREKYMALSLQSFCPTTRRYLQQLAEKPLETRTYEQGPDTPVSADAPVHPPALEQHPYEHCEPSTM 197
            ************************************************************************************************************************

hAMPD2-2    PGDLGLGLRMVRGVVHVYTRREPDEHCSEVELPYPDLQEFVADVNVLMALIINGPIKSFCYRRLQYLSSKFQMHVLLNEMKELAAQKKVPHRDFYNIRKVDTHIHASSCMNQKHLLRFIK 354
hAMPD2-4    PGDLGLGLRMVRGVVHVYTRREPDEHCSEVELPYPDLQEFVADVNVLMALIINGPIKSFCYRRLQYLSSKFQMHVLLNEMKELAAQKKVPHRDFYNIRKVDTHIHASSCMNQKHLLRFIK 360
hAMPD2-5    PGDLGLGLRMVRGVVHVYTRREPDEHCSEVELPYPDLQEFVADVNVLMALIINGPIKSFCYRRLQYLSSKFQMHVLLNEMKELAAQKKVPHRDFYNIRKVDTHIHASSCMNQKHLLRFIK 317
            ************************************************************************************************************************

hAMPD2-2    RAMKRHLEEIVHVEQGREQTLREVFESMNLTAYDLSVDTLDVHADRNTFHRFDKFNAKYNPIGESVLREIFIKTDNRVSGKYFAHIIKEVMSDLEESKYQNAELRLSIYGRSRDEWDKLA 474
hAMPD2-4    RAMKRHLEEIVHVEQGREQTLREVFESMNLTAYDLSVDTLDVHADRNTFHRFDKFNAKYNPIGESVLREIFIKTDNRVSGKYFAHIIKEVMSDLEESKYQNAELRLSIYGRSRDEWDKLA 480
hAMPD2-5    RAMKRHLEEIVHVEQGREQTLREVFESMNLTAYDLSVDTLDVHADRNTFHRFDKFNAKYNPIGESVLREIFIKTDNRVSGKYFAHIIKEVMSDLEESKYQNAELRLSIYGRSRDEWDKLA 437
            ************************************************************************************************************************

hAMPD2-2    RWAVMHRVHSPNVRWLVQVPRLFDVYRTKGQLANFQEMLENIFLPLFEATVHPASHPELHLFLEHVDGFDSVDDESKPENHVFNLESPLPEAWVEEDNPPYAYYLYYTFANMAMLNHLRR 594
hAMPD2-4    RWAVMHRVHSPNVRWLVQVPRLFDVYRTKGQLANFQEMLENIFLPLFEATVHPASHPELHLFLEHVDGFDSVDDESKPENHVFNLESPLPEAWVEEDNPPYAYYLYYTFANMAMLNHLRR 600
hAMPD2-5    RWAVMHRVHSPNVRWLVQVPRLFDVYRTKGQLANFQEMLENIFLPLFEATVHPASHPELHLFLEHVDGFDSVDDESKPENHVFNLESPLPEAWVEEDNPPYAYYLYYTFANMAMLNHLRR 557
            ************************************************************************************************************************

hAMPD2-2    QRGFHTFVLRPHCGEAGPIHHLVSAFMLAENISHGLLLRKAPVLQYLYYLAQIGIAMSPLSNNSLFLSYHRNPLPEYLSRGLMVSLSTDDPLQFHFTKEPLMEEYSIATQVWKLSSCDMC 714
hAMPD2-4    QRGFHTFVLRPHCGEAGPIHHLVSAFMLAENISHGLLLRKAPVLQYLYYLAQIGIAMSPLSNNSLFLSYHRNPLPEYLSRGLMVSLSTDDPLQFHFTKEPLMEEYSIATQVWKLSSCDMC 720
hAMPD2-5    QRGFHTFVLRPHCGEAGPIHHLVSAFMLAENISHGLLLRKAPVLQYLYYLAQIGIAMSPLSNNSLFLSYHRNPLPEYLSRGLMVSLSTDDPLQFHFTKEPLMEEYSIATQVWKLSSCDMC 677
            ************************************************************************************************************************

hAMPD2-2    ELARNSVLMSGFSHKVKSHWLGPNYTKEGPEGNDIRRTNVPDIRVGYRYETLCQELALITQAVQSEMLETIPEEAGITMSPGPQ        798
hAMPD2-4    ELARNSVLMSGFSHKVKSHWLGPNYTKEGPEGNDIRRTNVPDIRVGYRYETLCQELALITQAVQSEMLETIPEEAGITMSPGPQ        804
hAMPD2-5    ELARNSVLMSGFSHKVKSHWLGPNYTKEGPEGNDIRRTNVPDIRVGYRYETLCQELALITQAVQSEMLETIPEEAGITMSPGPQ        761
            ************************************************************************************
```

**Figure S1. Sequence alignment of full length hAMPD isoforms and isozymes.** Catalytic domain is underlined. **A**, alignment of hAMPD 1, 2-2 and 3-1 isoforms. Active-site residues are highlighted in cyan and conserved residues are red. **B**, alignment of hAMPD2-2, 2-4 and 2-5 isozymes. Conserved catalytic residues are highlighted in yellow.

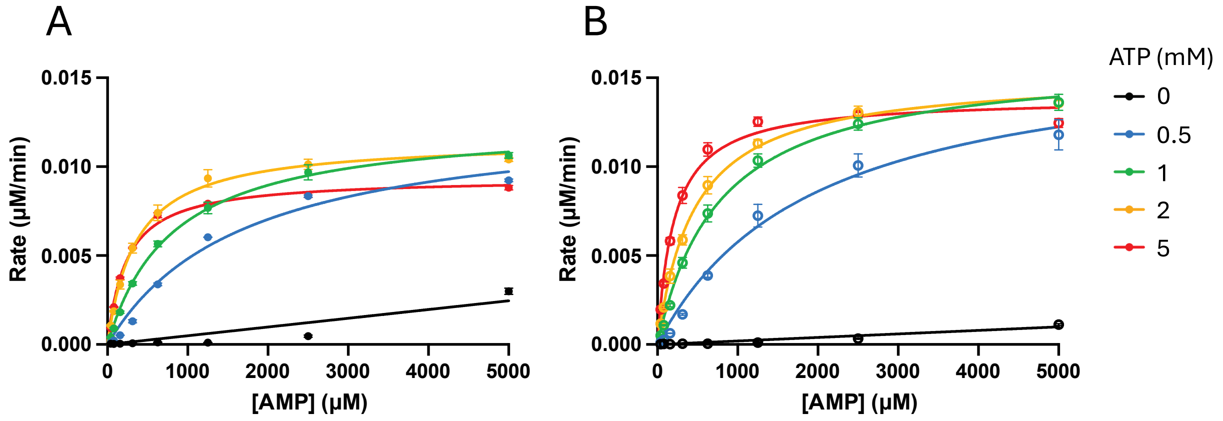


**Figure S2. ATP activates hAMPD2-2 and hAMPD2-2$^{\Delta128}$.** Panel A, full-length hAMPD2-2 (50 nM). Panel B, catalytic domain of hAMPD2-2 (5 nM).

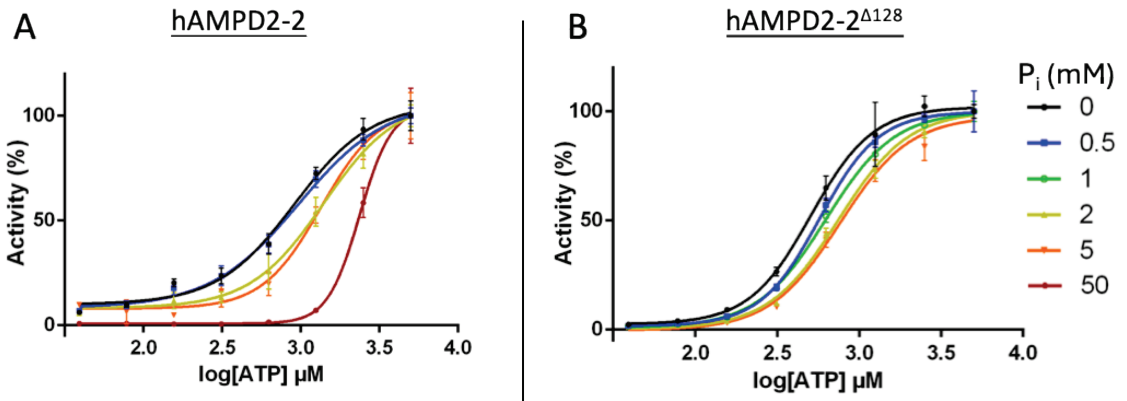


A

| $[P_i]$ (mM) | [ATP] at 50% activity (mM) |
|---|---|
| 0 | 0.91 |
| 0.5 | 0.97 |
| 2 | 1.4 |
| 5 | 1.3 |
| 50 | 2.3 |

B

| $[P_i]$ (mM) | [ATP] at 50% activity (mM) |
|---|---|
| 0 | 0.51 |
| 0.5 | 0.57 |
| 1 | 0.62 |
| 2 | 0.74 |
| 5 | 0.76 |

**Figure S3. Addition of $P_i$ with AMP-saturated full-length (Panel A) and catalytic domain of hAMPD2-2 (Panel B) under varying [ATP].** Activity of hAMPD2-2 was plotted varying [ATP] from 0 to 5 mM, with 1 mM AMP and increasing amounts of $P_i$. Data points were collected in triplicate, normalized to hAMPD2-2 activity at [ATP] = 5 mM for each $P_i$ concentration. Results were plotted using a 4-parameter fit curve in GraphPad prism, error bars represent standard deviations.

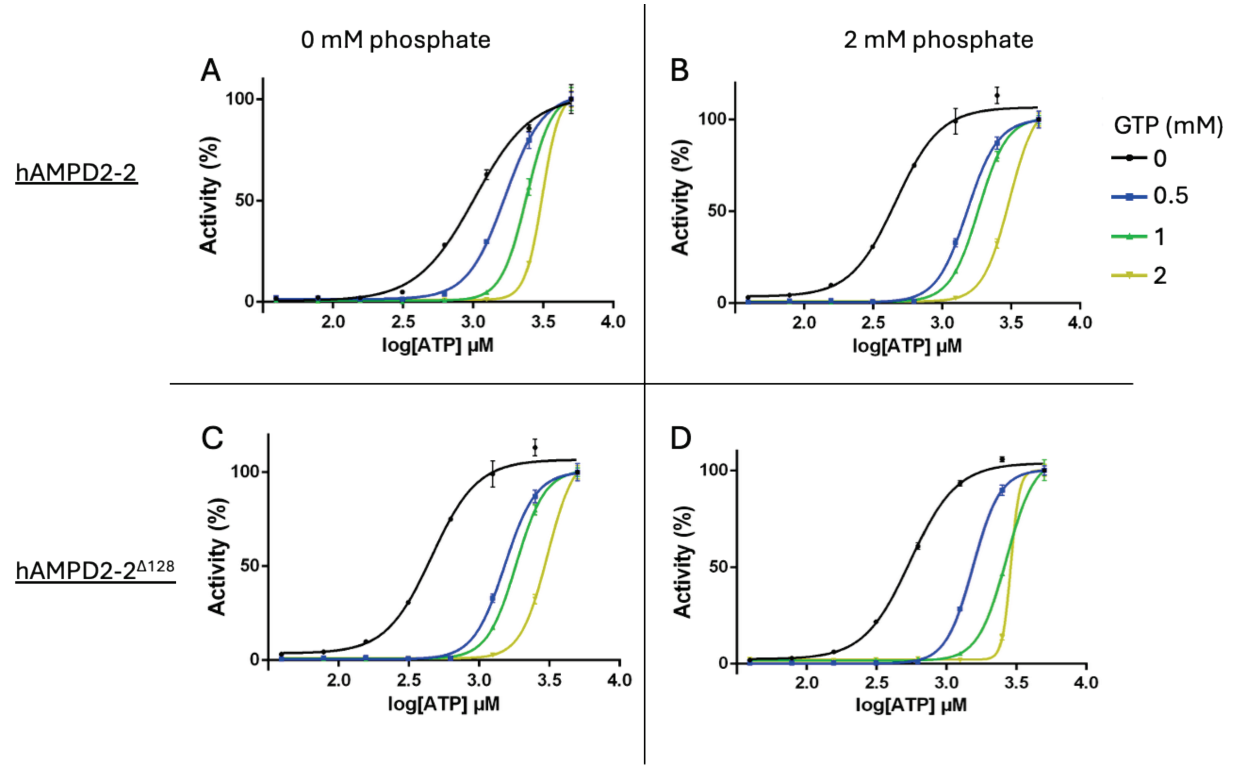


**Figure S4. Hill plots of substrate saturated hAMPD2-2 in the absence and presence of 2 mM $P_i$ with increasing concentrations of ATP, varying [GTP]**. Data was fitted to a four-parameter Hill plot. Results from A and B, full-length and C and D, catalytic domain of hAMPD2-2. Plots A and C are in the absence of $P_i$, B and D are in the presence of 2 mM $P_i$.

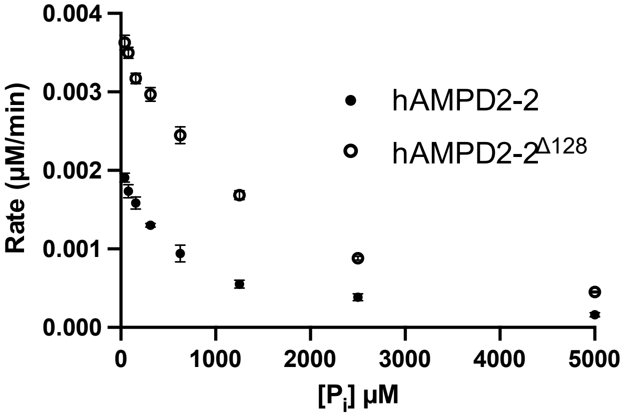


**Figure S5. Effect of GTP on saturated and activated hAMPD2-2, at increasing levels of $P_i$.** Data of full-length (50 nM, closed circles) and catalytic domain (5 nM, open circles) was determined at 1 mM AMP, 2 mM ATP, 1 mM GTP.

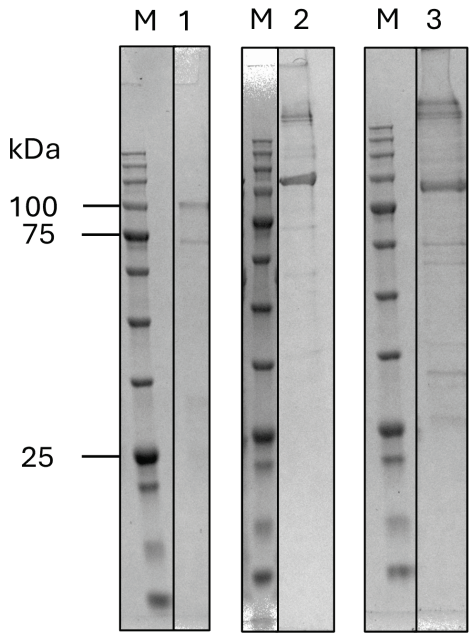


**Figure S6. Coomassie stained 12% SDS-Page of full-length and catalytic domain constructs of hAMPD2-2 purification from *E. coli***. M: molecular weight ladder (BLUEstain™ 2 Protein ladder, GoldBio), 1: $His_6$-hAMPD2-2, 94.4 kDa, 2: $His_6$-SUMO-hAMPD2-$2^{\Delta 128}$, 105.6 kDa, 3: $His_6$-SUMO-hAMPD2-$2^{\Delta 128}$, 91.3 kDa.

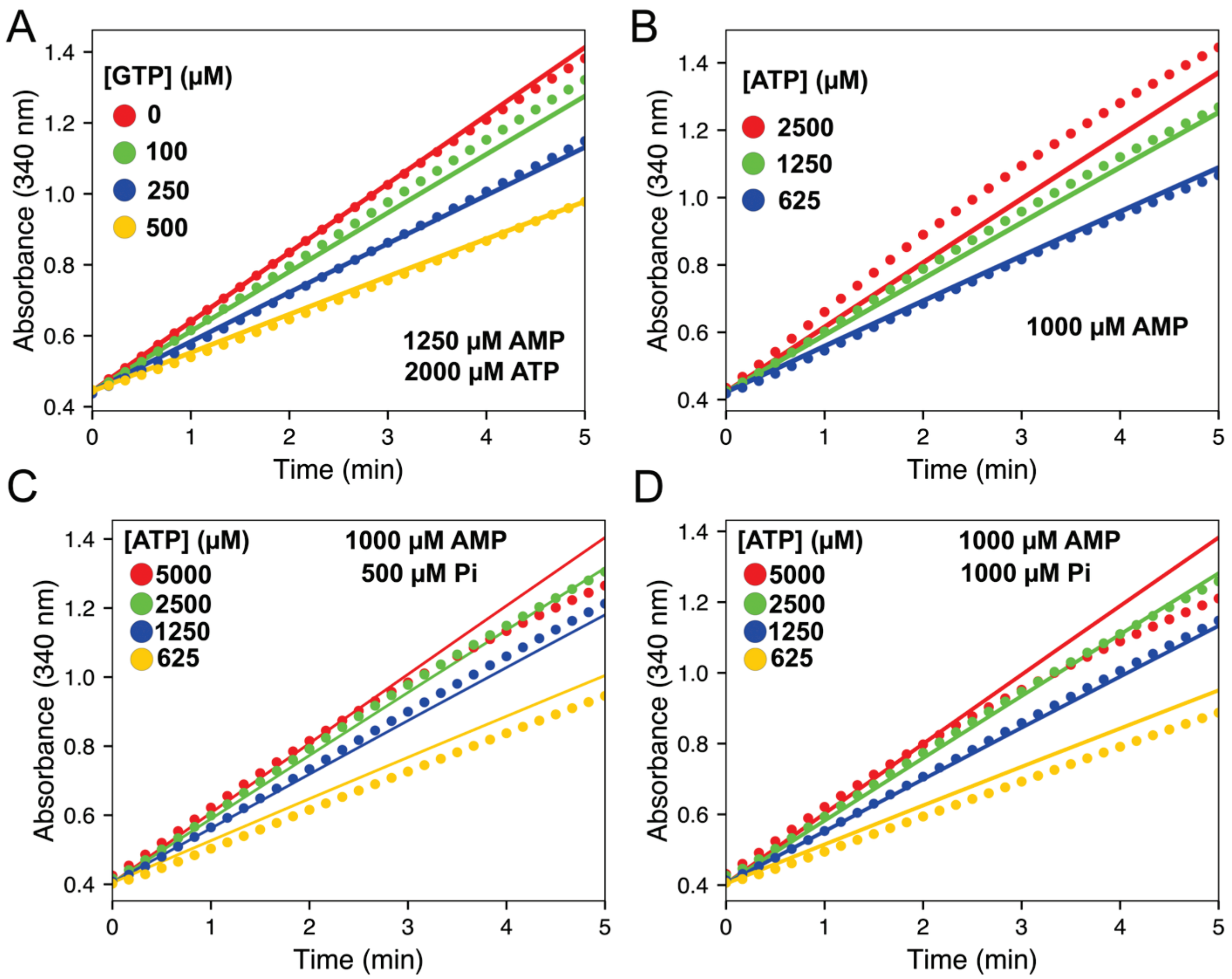


**Figure S7. Reduced global fitting of $A_{340}$ traces across GTP, ATP and $P_i$ concentrations.** Experimental absorbance traces at 340 nm are shown as colored circles, and the corresponding traces simulated by the kinetic model are shown as solid lines. (A) GTP titration at fixed 1250 μM AMP and 2000 μM ATP. (B) ATP titration at fixed 1000 μM AMP in the absence of added $P_i$. (C) ATP titration at fixed 1000 μM AMP in the presence of 500 μM $P_i$. (D) ATP titration at fixed 1000 μM AMP in the presence of 1000 μM $P_i$.

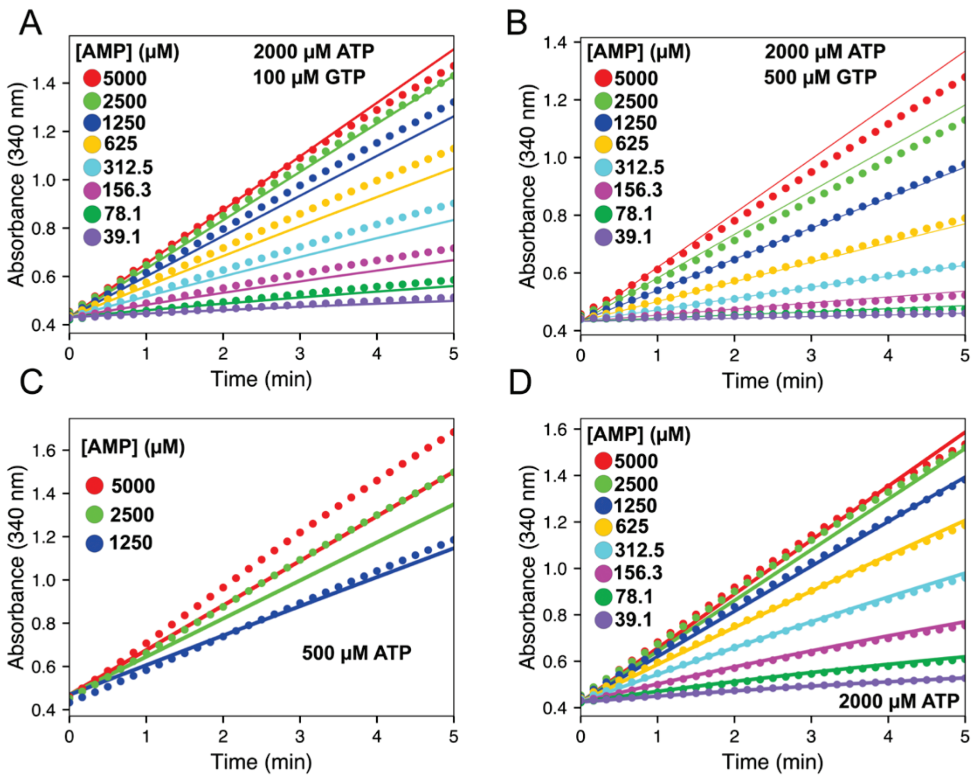


**Figure S8. Reduced global fits of AMP titrations at varying ATP and GTP conditions**. (A) AMP titration at fixed 2000 μM ATP in the presence of 100 μM GTP. (B) AMP titration at fixed 2000 μM ATP in the presence of 500 μM GTP. (C) AMP titration at fixed 500 μM ATP. (D) AMP titration at fixed 2000 μM ATP in the absence of added GTP.

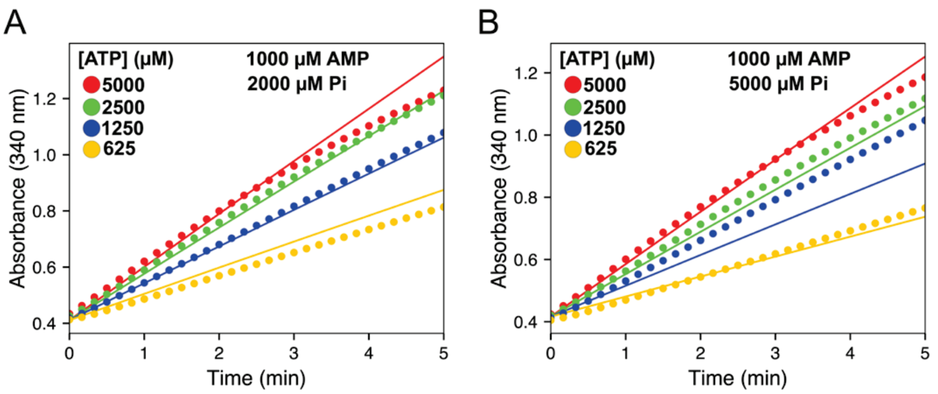


**Figure S9. Global fitting of ATP titrations at elevated $P_i$ concentrations.** Experimental absorbance traces at 340 nm are shown as colored circles, and the corresponding traces simulated by the reduced kinetic model are shown as solid lines. (A) ATP titration at fixed 1000 μM AMP and 2000 μM $P_i$. (B) ATP titration at fixed 1000 μM AMP and 5000 μM $P_i$.

### Supplementary Tables

**Table S1. Steady-state kinetic constants for activated enzyme with increasing $P_i$ concentrations.** Values are the mean determinations ± the standard deviation from three replicates. Results for hAMPD2-2 at 5 mM $P_i$ and above are from an enzyme preparation different from that used to determine values at concentrations below 2 mM $P_i$. Values determined using the data from Figures 4C and 4D.

| | $K_M$ (μM) | | $k_{cat}$ ($s^{-1}$) | | $k_{cat}/K_M$ ($M^{-1}s^{-1}$) | |
|---|---|---|---|---|---|---|
| [$P_i$] (mM) | hAMPD2-2 | hAMPD2-2$^{\Delta128}$ | hAMPD2-2 | hAMPD2-2$^{\Delta128}$ | hAMPD2-2 | hAMPD2-2$^{\Delta128}$ |
| 0 | 350 ± 20 | 340 ± 20 | 12.3 ± 0.1 | 140 ± 1 | (36 ± 12)x10$^3$ | (41 ± 2)x10$^4$ |
| 0.5 | 390 ± 10 | 380 ± 10 | 12.3 ± 0.1 | 135 ± 1 | (315 ± 8)x10$^2$ | (360 ± 10)x10$^3$ |
| 1 | 400 ± 20 | 380 ± 10 | 12.2 ± 0.1 | 132 ± 1 | (30 ± 2)x10$^3$ | (350 ± 10)x10$^3$ |
| 2 | 460 ± 20 | 410 ± 10 | 12.0 ± 0.1 | 129 ± 1 | (26 ± 1)x10$^3$ | (313 ± 8)x10$^3$ |
| 5 | 1300 ± 100 | 520 ± 20 | 39.4 ± 0.6 | 125 ± 4 | (31 ± 2)x10$^3$ | (24 ± 1)x10$^4$ |
| 10 | 430 ± 40 | 700 ± 40 | 28.9 ± 0.4 | 117 ± 1 | (67 ± 2)x10$^3$ | (170 ± 10)x10$^3$ |
| 20 | 900 ± 100 | 1300 ± 100 | 28.5 ± 0.5 | 119 ± 4 | (30 ± 4)x10$^3$ | (31 ± 8)x10$^3$ |
| 50 | 5000 ± 1000 | 9000 ± 2000 | 36 ± 2 | 119 ± 4 | (7 ± 1)x10$^3$ | (31 ± 3)x10$^3$ |

**Table S2. Titration of GTP with AMP saturated and ATP activated hAMPD2-2 in the absence and presence of $P_i$.** Enzyme rates (mol/min) of hAMPD2-2 and hAMPD2-2$^{\Delta128}$ at 2.5 mM ATP under saturating AMP concentration (1 mM), with 0- and 2-mM $P_i$, under varying GTP concentrations.

| [GTP] (mM) | hAMPD2-2 | | hAMPD2-2$^{\Delta128}$ | |
|---|---|---|---|---|
| | 0 mM $P_i$ | 2 mM $P_i$ | 0 mM $P_i$ | 2 mM $P_i$ |
| 0 | 1.8 ± 0.03 | 1.8 ± 0.03 | 8.1 ± 0.4 | 7.8 ± 0.1 |
| 0.5 | 1.7 ± 0.09 | 1.1 ± 0.03 | 6.9 ± 0.3 | 6.3 ± 0.2 |
| 1 | 1.1 ± 0.08 | 0.34 ± 0.01 | 5.8 ± 0.2 | 2.8 ± 0.04 |
| 2 | 0.29 ± 0.01 | 0.043 ± 0.004 | 1.9 ± 0.2 | 0.30 ± 0.03 |

**Table S3. Kintek global fitting model values.** Best-fit values, standard errors, units, and fitting status are shown listed for all reactions in the Kintek global fitting model. Abbreviations: E, enzyme; S, AMP; A, ATP; G, GTP; F, inorganic phosphate; P, product; Fa, allosteric phosphate-bound state; Fs, active-site phosphate-bound state. Thus, EA, ES, ESA, EG, and EAG denote nucleotide-bound enzyme states, whereas EFa, ESFa, and EGFa denote allosteric phosphate-bound states and EFs and EAFs denote active-site phosphate-bound states. Gray shading denotes parameters fixed during fitting. Red shading denotes poorly constrained fitted parameters with large uncertainties or non-determined errors.

| | Reaction | Parameter | Best-fit | Standard Error | Units | Status |
|---|---|---|---|---|---|---|
| 1 | E + S ⇌ ES | k+1 | 0.000142 | 0.008775 | $\mu M^{-1} s^{-1}$ | fitted |
| 1 | E + S ⇌ ES | k-1 | 772.01 | — | $s^{-1}$ | fitted |
| 2 | E + A ⇌ EA | k+2 | 0.16667 | — | $\mu M^{-1} s^{-1}$ | fixed |
| 2 | E + A ⇌ EA | k-2 | 701.37 | 224.85 | $s^{-1}$ | fitted |
| 3 | ES + A ⇌ ESA | k+3 | 0.16667 | — | $\mu M^{-1} s^{-1}$ | fixed |
| 3 | ES + A ⇌ ESA | k-3 | 0.010683 | 2.7744 | $s^{-1}$ | fitted |
| 4 | EA + S ⇌ ESA | k+4 | 5.9745 | 1.3485 | $\mu M^{-1} s^{-1}$ | fitted |
| 4 | EA + S ⇌ ESA | k-4 | 494.69 | 180.74 | $s^{-1}$ | fitted |
| 5 | ESA ⇌ EA + P | k+5 | 132.94 | 1.7346 | $s^{-1}$ | fitted |
| 5 | ESA ⇌ EA + P | k-5 | 0.016667 | — | $\mu M^{-1} s^{-1}$ | fixed |
| 6 | E + G ⇌ EG | k+6 | 0.10733 | — | $\mu M^{-1} s^{-1}$ | fixed |
| 6 | E + G ⇌ EG | k-6 | 29.426 | 7.6008 | $s^{-1}$ | fitted |
| 7 | EA + G ⇌ EAG | k+7 | 0.16667 | — | $\mu M^{-1} s^{-1}$ | fixed |
| 7 | EA + G ⇌ EAG | k-7 | 10.135 | 2.0936 | $s^{-1}$ | fitted |
| 8 | E + F ⇌ EFa | k+8 | 0.016667 | — | $\mu M^{-1} s^{-1}$ | fixed |
| 8 | E + F ⇌ EFa | k-8 | 1.37e+04 | 1.6e+04 | $s^{-1}$ | fitted |
| 9 | ES + F ⇌ ESFa | k+9 | 0.16667 | — | $\mu M^{-1} s^{-1}$ | fixed |
| 9 | ES + F ⇌ ESFa | k-9 | 0.034917 | 15.581 | $s^{-1}$ | fitted |
| 10 | EFa + S ⇌ ESFa | k+10 | 2.2419 | 8.6559 | $\mu M^{-1} s^{-1}$ | fitted |
| 10 | EFa + S ⇌ ESFa | k-10 | 3.1107 | 28.322 | $s^{-1}$ | fitted |
| 11 | EG + F ⇌ EGFa | k+11 | 0.16667 | — | $\mu M^{-1} s^{-1}$ | fixed |
| 11 | EG + F ⇌ EGFa | k-11 | 3.33e+04 | 5.62e+07 | $s^{-1}$ | fitted |
| 12 | EFa + G ⇌ EGFa | k+12 | 0.16667 | — | $\mu M^{-1} s^{-1}$ | fixed |
| 12 | EFa + G ⇌ EGFa | k-12 | 11.133 | 9.56e+05 | $s^{-1}$ | fitted |
| 13 | E + F ⇌ EFs | k+13 | 0.16667 | — | $\mu M^{-1} s^{-1}$ | fixed |
| 13 | E + F ⇌ EFs | k-13 | 1.75e+03 | 1.56e+04 | $s^{-1}$ | fitted |
| 14 | EA + F ⇌ EAFs | k+14 | 0.16667 | — | $\mu M^{-1} s^{-1}$ | fixed |
| 14 | EA + F ⇌ EAFs | k-14 | 7.78e+03 | 1.01e+05 | $s^{-1}$ | fitted |
| 15 | EFs + A ⇌ EAFs | k+15 | 0.019804 | 4.1704 | $\mu M^{-1} s^{-1}$ | fitted |
| 15 | EFs + A ⇌ EAFs | k-15 | 369.51 | 7.82e+04 | $s^{-1}$ | fitted |
| 16 | ESFa ⇌ EFa + P | k+16 | 2.2278 | 22.005 | $s^{-1}$ | fitted |
| 16 | ESFa ⇌ EFa + P | k-16 | 0.016667 | — | $\mu M^{-1} s^{-1}$ | fixed |
